# Modular cyanobacterial regulatory architecture enables dynamic graded responses to oxidative stress

**DOI:** 10.1101/2025.07.21.665879

**Authors:** Zachary Johnson, Natalie C. Sadler, Marci Garcia, Ashley Gilliam, Eric Allen Hill, Wei-Jun Qian, Margaret S. Cheung, Pavlo Bohutskyi

## Abstract

A fundamental paradox of oxygenic photosynthesis: growth-essential energy machinery generates reactive oxygen species (ROS) threatening survival, yet the systems-level regulatory networks balancing the growth-survival trade-off remain unclear. Through integrative experimental-computational analysis combining steady-state transcriptomics with independent component analysis across 0-78.4% oxygen, we decoded the regulatory architecture driving progressive transitions in *Synechococcus elongatus* PCC 7942 from ROS sensing through defense to growth shutdown. Integration of 407 transcriptome samples identified 78 regulatory modules (iModulons) explaining 72.3% of expression variance and revealed calibrated responses: low stress triggers metalloregulators (SufR, PerR) for ROS sensing and primary antioxidants; moderate stress activates RpaB operating through four distinct regulatory states redirecting metabolism to defense; severe stress induces growth arrest via stringent response pathway convergence. This quantitative regulatory framework enables precise growth-defense calibration through modular network architecture: RpaB coordinates genome-wide resource reallocation (RpaB∼P growth-promoting, RpaB ROS defense-activating, RpaABC circadian-integrating, RpaB ycf46 checkpoint activation), offering systematic strategies for engineering stress-tolerant bioplatforms and predictive models for environmental stress responses.

## 1. Introduction

Photosynthetic organisms face a fundamental metabolic paradox: oxygenic photosynthesis simultaneously drives growth and generates reactive oxygen species (ROS) that impair cellular functions[1, 2]. This ancient challenge now critically limits cyanobacterial biotechnology, where oxidative stress constrains bioproduction when maximizing metabolic output[3, 4]. Decoding the systems-level regulatory networks that orchestrate cyanobacterial oxidative stress responses is therefore essential for both understanding emergent properties of cellular stress adaptation and engineering predictable stress-tolerant bioplatforms.

The cellular challenge is formidable. ROS primarily originates from photosynthetic electron transport chain, where light-driven reactions generate superoxide and hydrogen peroxide[5, 6]. Environmental fluctuations in light and nutrients constantly perturb cellular redox balance[7–9]. In response, cyanobacteria have evolved sophisticated redox-sensing networks dynamically monitoring conditions and coordinating adaptive transcriptional responses[10, 11] involving hierarchical regulatory modules with global and pathway-specific regulators precisely allocating resources between growth and protection[7–9, 12].

A critical knowledge gap persists. While individual redox-sensitive regulators (SufR, PerR) and stress response regulators (RpaB[13, 14] are identified, are identified, the network-level interactions and emergent regulatory logic that coordinate graded graded responses across stress intensities—precisely calibrating defense activation with growth maintenance—remains unknown. This lack of systems-level integration prevents quantitative prediction and rational optimization of cyanobacterial performance under variable conditions. Without this integrated network understanding, engineering efforts remain trial-and-error, and natural population responses to increasing environmental stressors cannot be predicted.

*Synechococcus elongatus* PCC 7942 presents an exceptional opportunity to address these gaps. Its role in defining KaiABC circadian oscillator[15] and associated regulators RpaA[16], RpaB[13, 14], and sigma factors[17] has generated extensive transcriptomic datasets, enabling sophisticated data-driven analyses of circadian regulatory dynamics driven by changing redox environment [18, 19]. Key transcriptional regulators—RpaB, RpaA and Rre1—respond to redox signals[15, 20, 21], yet the quantitative network dynamics and modular organization governing their collective coordination across different stress intensities remains uncharacterized.

We developed an integrative experimental-computational framework combining steady-state physiology with systems-level transcriptomics to address these fundamental questions. By coupling turbidostat cultivation with systematic oxygen manipulation (0-374% air saturation), we maintained cells at steady state while quantifying both physiological and transcriptional responses across the complete stress continuum—from minimal stress through complete growth inhibition. This controlled perturbation approach enabled systematic mapping of network state transitions and quantitative correlation of regulatory module activities with physiological outcomes

Transcriptome samples and physiological measurements from the oxidative stress experiments in this study were integrated with comprehensive transcriptome profiles from public databases for PCC 7942 and a set of previously unpublished profiles collected by our team, creating an unprecedented dataset of 407 high-quality samples. We then applied independent component analysis to identify regulatory modules (iModulons)[22, 23]. Compared to alternative clustering methods, ICA has demonstrated remarkable results in reconstructing known regulatory interactions, making it ideal for decomposing complex multi-layered regulatory networks with multiple active regulators. Recent ICA analysis of PCC 7942 has provided valuable insights into circadian regulation[18]. Here, we leverage this computational approach to systematically decode tiered oxidative stress responses, revealing stress-responsive regulatory modules and establishing quantitative correlations between iModulon activities and stress intensities and growth rates while discovering distinct network transition states for the first time.

Our analysis reveals sophisticated regulatory architecture where cells progress from specialized ROS sensing and defense through coordinated metabolic rewiring to systematic growth shutdown. Most remarkably, we uncover previously unrecognized connections to bacterial stringent response pathways, revealing network-level regulatory mechanisms with profound implications for both systems-level understanding and rational bioengineering. This work transforms our understanding of oxidative stress from a simple damage-response paradigm to a quantitative regulatory program balancing growth and survival.

## 2. Methods

### 2.1 Cyanobacteria Culture

*Synechococcus elongatus* PCC 7942 CscB/SPS strain, engineered to overexpress sucrose phosphate synthase (SPS) and sucrose permease (CscB), was obtained from Dr. Daniel Ducat [24–26]. Cultures were maintained in BG11 medium supplemented with chloramphenicol (12.5 mg L⁻¹) and kanamycin (25 mg L⁻¹). Standard growth conditions were maintained at 29±2°C with nitrogen gas sparging supplemented with 2% CO₂ under 200 μmol photons m⁻² s⁻¹ illumination from LED lights. Culture mixing was facilitated using PFTE Octagon Spinbar Magnetic Stirring Bars (9.5 mm x 25.4 mm) at 200 r.p.m. Antibiotics were removed prior to experimental procedures.

### 2.2 Photobioreactor Operation and Continuous Culture

Continuous cultivation was performed in 6.5-L New Brunswick Bioflo 310 fermentors (Eppendorf, Inc.) equipped with custom LED photobioreactors[27]. Steady-state cultures (5.5-L) were maintained in turbidostat mode at OD₇₃₀ of 0.08, with constant conditions of 250 rpm agitation, 30°C, and pH 8.0 (controlled by 2 M NaOH or HCl addition). Illumination was provided at 760 μmol photons m⁻² s⁻¹ using 630 and 680 nm narrow-band LED illuminator chips (Marubeni America Corporation). Photosynthetically active radiation (PAR, 400-700 nm) was monitored using three quantum sensors (LI-210SA, LI-COR Biosciences). Dissolved O₂ was measured using a Clark-type electrode (InPro® 6800Series, Mettler Toledo). Turbidostat operation was maintained by correlating transmitted irradiance measurements with spectrophotometer-verified OD_750_ values to control medium addition and removal.

### 2.3 Oxidative Stress Implementation

Initial conditions utilized a baseline sparging rate of 2.04 L min⁻¹ with 98:2 N_2_/CO_2_ gas mixture. Oxidative stress was implemented through stepwise increases in O_2_ content while maintaining constant total gas flow and CO_2_ concentration. O_2_ levels were increased sequentially to 5.12%, 9.8%, 19.6%, 29.4%, 39.2%, 49%, 58.8%, 68.6%, and 78.4%, with corresponding N₂ reduction. Steady-state conditions were confirmed by stable OD₇₃₀, pH, and dissolved O₂ measurements (≤3% variation) for each oxygen level. Growth rates were calculated from spent medium accumulation rates. Complete growth inhibition occurred after ∼40 hours exposure at 78.4% O₂ (∼375% air saturation). Samples for transcriptome analysis were collected after minimum three residence times at steady-state or ∼50 hours for 78.4% O₂. The complete experimental series spanned approximately 40 days. Air saturation percentages were calculated according to:

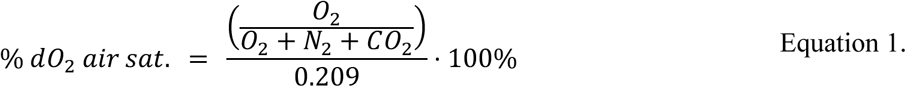

Growth rate dose-response modeling was performed to characterize stress response patterns and quantify physiological transition points. Multiple mathematical models were evaluated for goodness of fit to the experimental growth rate data, including a modified sigmoidal Hill-type equation, Gompertz-type inhibition model, modified power law inhibition, and classic Hill inhibition equation (equations provided in **Supplementary Methods 1**). Model parameters were estimated using nonlinear optimization by minimizing root mean square normalized deviations, and model performance was evaluated using the coefficient of determination (R²).

### 2.4 Microscopy and cell area quantification

Cells were stained with 2X SYBR Gold, dark-incubated for 20 minutes, then transferred to 12-well chambers (Ibidi) mounted on 75×24 mm #1.5 glass coverslips. After an additional 20-minute dark incubation, samples were imaged using a Leica inverted confocal microscope with 63X objective. Cell dimensions were analyzed from captured images using standard image analysis software.

### 2.7 RNA Isolation and Sequencing

Total RNA was extracted using RNeasy kit with QiaShredder columns (QIAGEN). Cells were lysed in RLT buffer containing 1% β-mercaptoethanol, homogenized through QiaShredder, then processed following manufacturer’s protocol with 70% ethanol addition. Two biological replicates were processed per condition. Samples underwent DNase treatment and rRNA depletion before paired-end sequencing (2×150bp, ∼350M reads, ∼105GB) with single indexing on Illumina platform at Azenta US, Inc.

### 2.8 RNA-seq Processing

RNA-seq reads were aligned and quantification with Rsubread[28]. The reference was collected from NCBI RefSeq November 16, 2022[29]. Reference replicons include the chromosome (NC007604.1), and plasmids pANL (NC_007595.1), pANS (NC_004990.1). Reference sequences were functionally annotated by integrating gene annotations from multiple annotation pipelines (**Supplementary Methods 2.1**). Literature references used in the annotation include circadian peak expression identified in Markson et al., 2013[16], and transcription start site and operons identified in Vijayan et al., 2011[30]. The regulatory network compiled identifies 701 potential regulatory interactions based on a combination of experimental and computational evidence. For further detail see **Supplementary Methods 2.2 and Supplementary Data 1**.

### 2.9 Independent Component Analysis of Gene Expression

In total, 557 RNA-seq samples representing 283 unique conditions from 20 unique RNA-sequencing projects were collected. Of these, 385 samples and 209 conditions were collected from literature while 172 samples and 74 conditions represent internally generated data sets. These data sets correspond to three large scale projects characterizing: coculture perturbations (96 samples, 41 conditions), circadian sucrose product (30 samples, 10 conditions), and oxidative stress perturbations 46 samples 23 conditions). In total, this dataset represents the most comprehensive collection of RNA-seq data for *S. elongatus PCC 7942* to date. Integration of diverse experimental conditions was essential for robust iModulon identification. A large number of diverse transcriptome profiles from experiments spanning circadian rhythms, nutrient limitations, genetic perturbations, various light intensities, and environmental stresses provided the statistical power and regulatory diversity necessary to distinguish genuine stress-responsive iModulons from experimental artifacts. Cross-condition validation confirmed that our oxidative stress iModulons represent authentic regulatory units rather than condition-specific noise. After removing outliers based on global correlation, replicate correlation < 0.90 and protein-coding reads below 5×10⁵ the final high-quality dataset comprised of 407 samples representing 73% of initial samples, all newly collected samples passing QC controls (**Supplementary Data 2**).

Expression data was transformed to log transcripts per million reads (logTPM) and normalized to project-specific reference condition. Optimal Independent component analysis (ICA)[23] was applied to transformed normalized counts resulting in identification of 78 iModulons at an optimal dimensionality of 160, explaining 73.41% of variance in the gene expression data. Regulatory iModulons were identified at q-value < 0.001 while functional iModulons were identified at a q-value < 0.01. In total, 43 iModulons were enriched for either regulator targets (31 iModulons) or functional gene sets (31 iModulons). For further details see **Supplementary Methods 3,** iModulon gene sets are given in **Supplementary Data 3.**

### 2.10 Explained variance of gene expression by independent components

To quantify the contribution of each independent component (iModulon) to the overall expression matrix, we computed the cumulative explained variance (CEV) as follows:

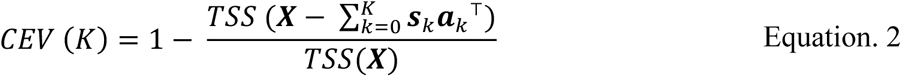

Here ***X*** ∈ ℝ*^m^* ^×^ *^n^* represents the original gene expression matrix with *m* genes (rows) and *n* samples (columns). Each component *k* is defined by an iModulon signal vector ***s**_k_* ∈ ℝ*^n^* (columns of signal matrix S) and a corresponding gene weight vector ***a**_k_* ∈ ℝ*^n^* (columns of mixing matrix A), where ***s**_k_**a**_k_*^T)^reconstructs the rank-1 approximation of the expression signal attributed to component *k*. The total sum of squares (TSS) is defined as:

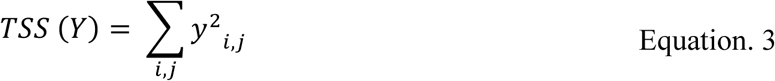

We compute CEV across all samples in ***X***, as well as for a subset of samples corresponding to the oxygen perturbation experiment, referred to as explained variance O₂ stress. Additionally, we calculate the proportion of variance explained for the subset of genes significantly weighted in each component, referred to as gene set explained variance. These significant genes are identified from the mixing vector ***a***_!_ using the D’Agostino K² threshold, described previously[31].

## 3. Results & Discussion

### Systems-Level Characterization of Escalating Oxidative Stress Reveals Progressive Physiological Transitions and Multi-Phase Regulatory Network States in *S. elongatus* PCC 7942

To elucidate the systems-level regulation governing cyanobacterial graded defense activation and growth maintenance across oxidative stress levels, we systematically characterized S. elongatus physiological responses to elevated O₂ under high light (760 μmol photons m⁻² s⁻¹). Using a turbidostat-based experimental system with precise control of dissolved oxygen (**Figure 1a**), we exposed steady-state cultures to incrementally increasing O₂ concentrations ranging from 0% to 78.4% (gas phase), corresponding to dissolved O₂ levels of up to 374% air saturation. Steady-state samples collected across this stress gradient were subjected to physiological and transcriptome analysis to determine key control points and the transcriptional architecture driving dynamic stress responses in *S. elongatus.* This approach uncovered a progressive response continuum comprising three distinct regimes, each characterized by specific growth and morphological adaptations (**Figure 1b**, **Table 1**).

**Figure 1.**
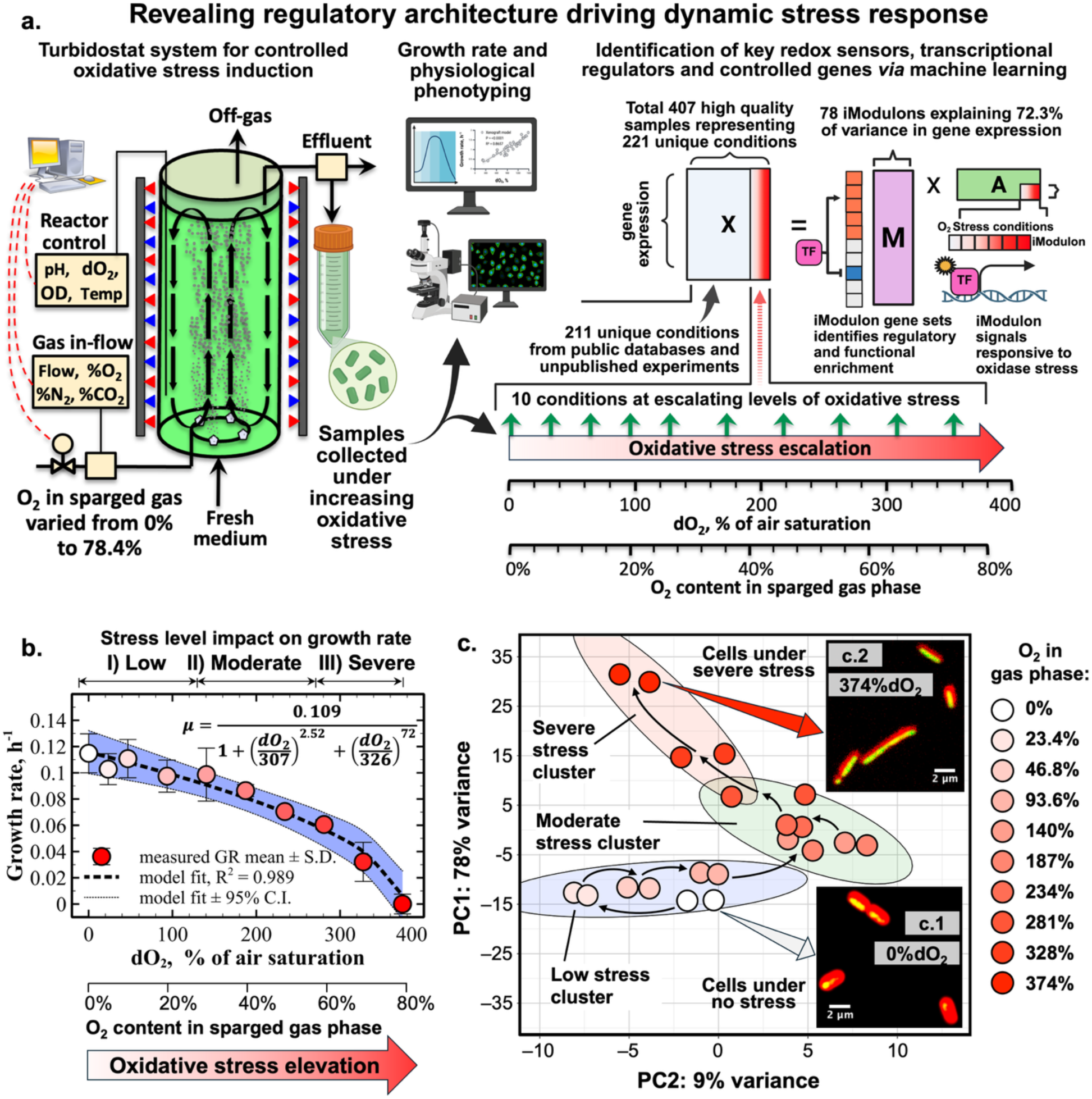
*Synechococcus elongatus* PCC 7942 exhibits a progressive physiological response to escalating oxidative stress with identifiable regulatory transition points. (**a)** Experimental workflow showing turbidostat photobioreactor system for controlled oxidative stress induction and downstream analytical approaches for physiological and transcriptional characterization. Transcriptomic profiles from this oxidative stress study were integrated with profiles from public databases and unpublished studies to create a comprehensive dataset of 407 high-quality samples describing 221 unique expression states for independent component analysis. **(b)** Growth rate exhibits continuous decline across the oxidative stress gradient, revealing three distinct physiological regimes. Steady-state growth rates (circles, mean ± SD) measured after complete acclimation show sigmoidal dose-response behavior fitted with modified Hill equation (dashed line, R² = 0.989). Model parameters identify IC₅₀ at 307% O₂ saturation, marking critical transition to severe stress response. Colored regions (95% confidence intervals) delineate analytical regimes: Low stress (0-140% O₂ saturation, 14% growth reduction), Moderate stress (140-280%, 47% cumulative reduction), and Severe stress (280-374%, complete growth arrest). **(c)** Principal component analysis (PCA) of steady-state gene expression profiles under varying levels of oxidative stress, demonstrates systematic progression through stress regimes. Insets (**c.1**, **c.2**) show representative cell morphology under lowest stress (0% O₂ air saturation) versus highest stress (374% O₂) conditions (chlorophyll autofluorescence, red; SYBR Gold DNA stain, green; scale bar = 2 μm).

**Table 1.** Progressive physiological transitions reveal dynamic stress inhibition and response regimes across the oxidative stress continuum. Major transitions in *S. elongatus* PCC 7942 response to oxidative stress distill into three analytical phases capturing graduated cellular progression from effective defense through metabolic reprogramming to coordinated survival programs.

| Stress regime | O<sub>2</sub> Content Range (% in gas phase) | Dissolved O<sub>2</sub> Range (as % of air saturation) | Key Characteristics |
| --- | --- | --- | --- |
| I. Low stress | ~0-30% | 0-140% | Growth rate minimal reduction: 14% from baseline<br>Progressive increase in cell size indicating activation of stress response<br>Stable physiological parameters<br>Effective cellular defense systems maintain near-normal growth despite oxidative stress exposure |
| II.<br>Moderate<br>stress | ~30-60% | 140-280% | • Growth rate reduction slope: 2.5-fold increase vs. Low stress regime<br>• Cumulative growth rate decrease: 47% from baseline<br>• Stable mean cell size with increased variability<br>• Defense systems continue supporting growth but with measurable metabolic cost, creating growth-protection trade-offs |
| III.<br>Severe<br>stress | ~60-80% | 280-375% | • Growth rate reduction slope: 2.8-fold increase vs. Moderate stress regime<br>• Terminal growth inhibition after ~40 hours at max O<sub>2</sub> level<br>• Dramatic changes in cell morphology and size<br>• Cellular systems shift from active defense to survival-oriented programs when stress exceeds adaptive capacity |

The low stress regime (∼0-30% O₂, 0-140% air saturation) featured minimal growth reduction (14% from baseline) despite a progressive increase in cell size, indicating activation of stress response mechanisms. During the moderate stress regime (∼30-60% O₂, 140-280% air saturation), we observed a 2.5-fold increase in the growth rate reduction slope compared to low stress, resulting in 47% decrease from baseline. Cells maintained relatively stable mean size but with increased variability. Under severe stress (∼60-80% O₂, 280-375% air saturation), the growth rate reduction slope increased another 2.8-fold compared to moderate regime, culminating in complete growth inhibition after approximately 40 hours at maximum O₂ levels, accompanied by dramatic changes in cell morphology.

Mathematical modeling validated distinct response patterns and quantified stress tolerance thresholds. A modified sigmoidal Hill-type equation (Figure 1b) best fit experimental data (R² = 0.989) versus alternative models: Gompertz-type inhibition[32], modified power law inhibition[33], and classic Hill inhibition equation[34] (**Supplementary Figure 1**). The modified Hill-type model predicted a maximum growth rate without inhibition of 0.109 h⁻¹, with an inhibitory concentration for 50% growth rate reduction (IC₅₀) at dO₂ of 307% of air saturation (64.3% O₂ in gas phase), identifying transition to severe stress conditions.

Principal component analysis (PCA) of the transcriptome data revealed systematic progression of cellular adaptation that corresponded closely to the three physiological regimes identified through growth rate analysis (**Figure 1c**). Hierarchical clustering analysis was consistent with these distinct transcriptional regimes **(Supplementary Figure 2).** The first principal component (PC1) explained 78% of variance, capturing the magnitude of transcriptional changes in response to increasing O₂ levels, while the second principal component (PC2), accounting for 9% of variance, reflected transitions between stress response regimes, particularly highlighting the distinct transcriptional state under severe stress conditions.

Confocal microscopy revealed dramatic changes in cellular morphology in response to oxidative stress. Quantitative analysis showed a >30% increase in mean cell area with significant reduction in cell width under high O₂ conditions compared to minimal oxidative stress (**Figure 1 c.1-c.2**). To systematically characterize how these morphological adaptations developed across the stress continuum, we tracked multiple cellular parameters including ash-free dry weight, size distribution through membrane filtration, and mean cell area based on chlorophyll autofluorescence at each oxygen concentration (**Figure 2a-b**). This detailed characterization revealed that morphological adaptations, including reduction in cell thickness and elongation, began during early stress exposure, well before significant growth inhibition occurred. Importantly, morphological trends matched the three response regimes identified through growth rate and transcriptomic analyses, providing independent physiological confirmation of the underlying regulatory transitions.

**Figure 2.**
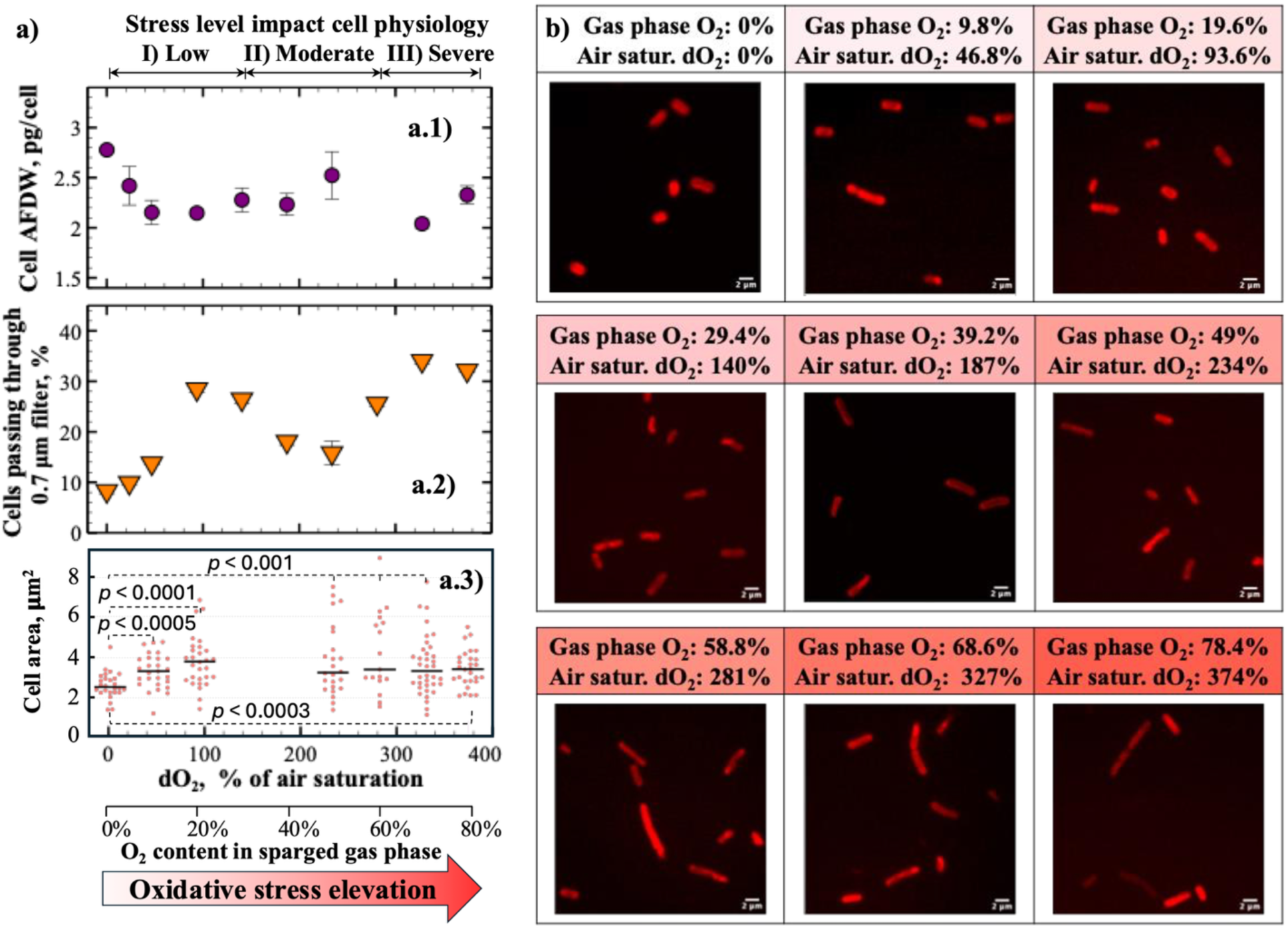
Progressive morphological adjustments track continuous stress response in *S. elongatus* PCC 7942. **(a)** Quantitative cellular metrics reveal gradual morphological transitions: **(a.1)** ash-free dry weight per cell, **(a.2)** size distribution shifts validated by filtration analysis using 0.7 μm membrane, **(a.3)** individual cell area measurements (n=30 cells per condition, horizontal bars indicate means). **(b)** Confocal microscopy confirms continuous morphological adaptation across the complete stress gradient (chlorophyll autofluorescence, red; scale bar = 2 μm), demonstrating coordinated physiological responses to escalating oxidative stress.

These distinct physiological regimes raised a fundamental question: what molecular mechanisms orchestrate such precise transitions between cellular states? Our transcriptomic analysis provided remarkable answers.

### Decoding Systems-Level Transcriptional Architecture Orchestrating Dynamic Stress Response Programs

The physiological response regimes immediately suggested underlying molecular machinery orchestrating these transitions. To decode these molecular mechanisms, we employed Independent Component Analysis (ICA), which decomposes transcriptomic data into independently regulated gene modules (iModulons). Our analysis identified 78 iModulons in PCC 7942 (**Supplementary Figure 4**), explaining 72.3% of variance across 407 RNA-seq samples and 65.1% in our oxidative stress dataset. Quantitative correlations between iModulon activities and both stress intensities and growth rates enabled direct linkage of transcriptional states to physiological outcomes across the complete oxidative stress continuum. The analysis identified previously unreported regulatory modules governing metal homeostasis and cofactor biosynthesis, revealed redox-sensing and multi-state response regulator networks, detailed photosynthetic and metabolic pathway organization, and demonstrated circadian-stress integration with stringent response-mediated shutdown under severe stress (**Supplementary Figure 5**).

Of the 78 iModulons, 41 show enrichment for known TF regulon targets (q-value < 1e^−3^) or functional gene sets (q-value < 0.01), with 31 defined by regulatory control and 12 by functional enrichment, enabling direct biological interpretation of stress-responsive gene regulation. Diverse RpaA and RpaB-related regulons emerged as the strongest contributors to global gene expression variance, highlighting their central role in oxidative stress response. Notably, our analysis revealed that RpaB operates through multiple independently regulated states rather than as a single regulon, while identifying previously uncharacterized regulatory modules controlling redox sensing, metal homeostasis, and antioxidant defense—providing a more nuanced regulatory landscape view.

Analysis of iModulon activities across the complete oxygen gradient revealed seven key molecular markers with strong correlations (|r| > 0.90) to oxidative stress: SufR (r = 0.99), RpaB ROS (r = 0.95), sigG rpoD2 (r = 0.94), PBS (r = -0.95), and RpaB∼P (r = -0.92) (**Figure 3a, b**). These iModulons effectively track stress progression from minimal to severe conditions. Crucially, when examining their relationship with growth rate, we discovered a striking inverse pattern: growth-promoting iModulons (RpaB∼P, r = 0.99, PBS, r = 0.84) showed strong positive correlation with growth rate, while stress-responsive iModulons (sigG rpoD2, r = -0.99; RpaB ROS, r = -0.90; SufR, r = -0.89) showed strong negative correlation. This inverse relationship directly links transcriptional states to physiological outcomes, demonstrating that oxidative stress systematically shifts cells from growth-promoting to survival-oriented gene expression programs. Examining iModulons across stress regimes revealed distinct activation patterns corresponding to our physiological observations (**Figure 3c**). During low stress (0-140% air saturation), early-response iModulons activated rapidly while growth remained largely unaffected. The SufR iModulon (r = 0.98) led this response, explaining exceptional 94.7% of its gene set variance and indicating tight regulatory control over iron-sulfur cluster assembly. Additional early-stress responders included Prophage (r = 0.98), REase (r = 0.94), and CCM-2 (r = 0.82), signaling immediate deployment of DNA repair and carbon concentration mechanisms.

**Figure 3.**
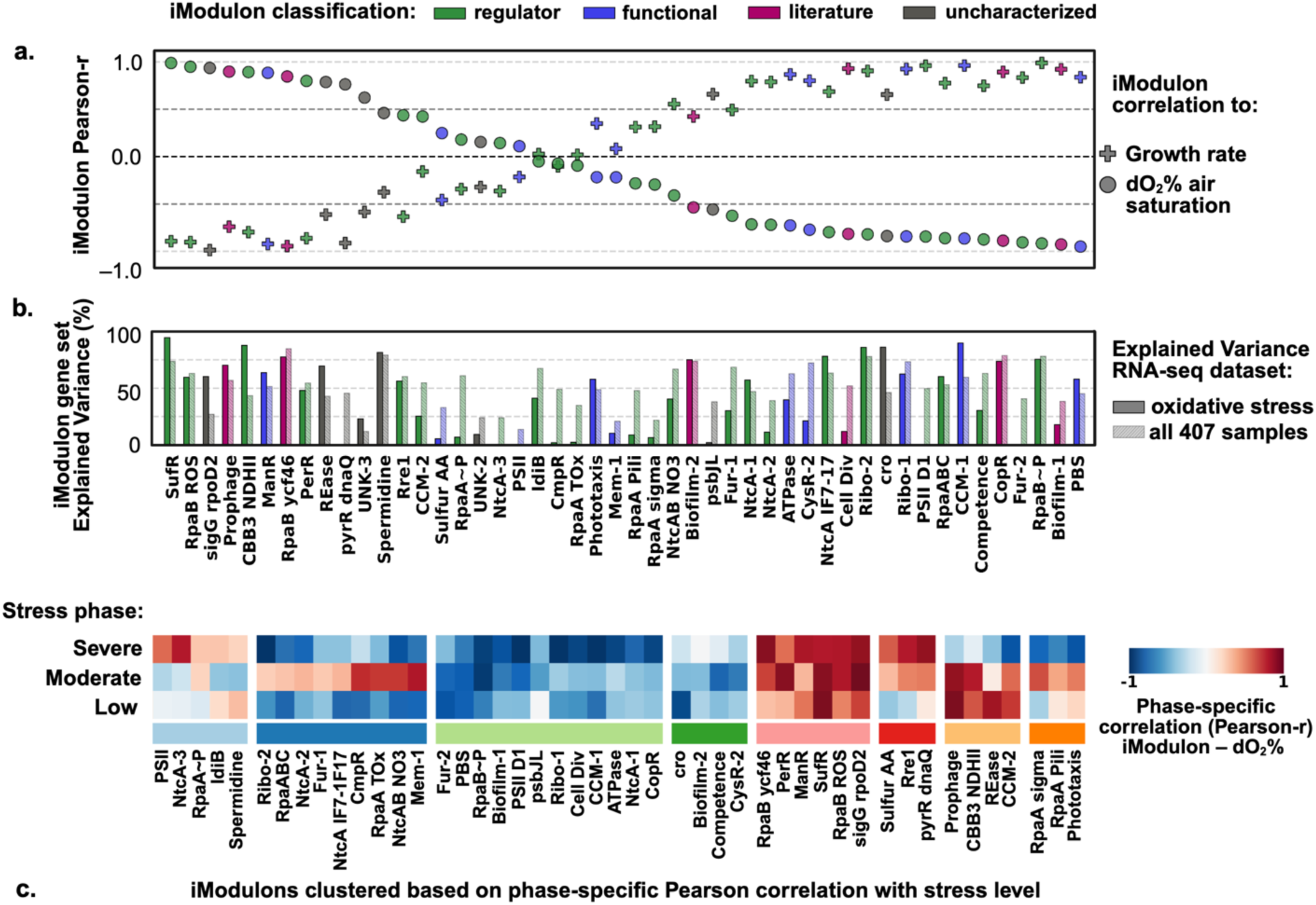
Independent component analysis reveals hierarchical stress-responsive iModulons with growth-correlated and stress-regime-specific activation patterns. **(a)** iModulons ranked by correlation across the complete oxidative stress gradient reveal hierarchical stress-responsive regulatory modules. Dual correlation analysis demonstrates relationships with both dissolved O₂ (circles) and growth rate (crosses) using all stress experiment samples. Primary stress sensors SufR (r = 0.99) and RpaB ROS (r = 0.95) show strongest O₂ correlations, while growth-promoting iModulons exhibit strong negative correlations: PBS (r = -0.95), RpaB∼P (r = -0.92), and CCM-1 (r = -0.86). Inverse correlation patterns demonstrate systematic transcriptional switching from growth to survival programs. **(b)** Gene set explained variance evaluates regulatory coordination within iModulons, identifying signals with the highest predictive power for each module gene set. Top stress-responsive iModulons explain 60-94% of constituent gene expression variance, confirming robust capture of gene expression signal. Colors indicate functional classification: regulatory (transcription factor target-enriched), functional (pathway-enriched), literature-defined, or uncharacterized modules. **(c)** Stress-regime-specific correlation analysis reveals diverse iModulon regulatory strategies: continuous dose-response relationships, biphasic activation with regime-specific peaks, threshold-activated responses, and switch-like state transitions. These distinct activation patterns reflect the progressive physiological response continuum observed in Figure 1, demonstrating how coordinated transcriptional programs orchestrate dynamic response to escalating stress.

In moderate stress (140-280% air saturation), the stress response expanded markedly. The PerR iModulon reached peak activation (r = 0.97), controlling hydrogen peroxide detoxification systems and explaining 48.1% of its gene set variance. Distinct iModulons activated specifically during moderate stress: Mem-1 (r = 0.91), CmpR (r = 0.84), and NtcAB NO_3_ (r = 0.82). Critically, several iModulons including CBB3 NDHII, Prophage, and CCM-2 showed biphasic behavior— strongly activated during low-to-moderate stress but plateaued or suppressed under severe stress— revealing a regulatory transition between active defense and metabolic shutdown.

Under severe stress (280-375% air saturation) a dramatic regulatory shift occurred, characterized by convergent growth-arrest programs. Newly activated iModulons included RpaB ycf46 (r = 0.97), pyrR dnaQ (r = 0.96), ManR (r = 0.92), and Rre1 (r = 0.92). This activation coincided with near-complete repression of growth-essential functions: photosynthesis (PSII D1, r = -0.99), carbon fixation (CCM-1, r = -0.99), protein synthesis (Ribo-1,2, r = -0.99), metal homeostasis (CopR, r = -0.96), cell division (Cell Div, r = -0.96), and ATP generation (ATPase, r = -0.96). The severity of this repression—with correlation coefficients approaching -1.0— indicates a coordinated cellular shutdown program activated under extreme oxidative stress.

To systematically investigate how these regulatory modules orchestrate responses across stress regimes, we prioritized iModulons based on three key criteria: correlation strength with specific stress regimes (|r| > 0.80), explained variance within their gene sets, and biological relevance to oxidative stress. This revealed distinct regulatory strategies for each regime. Under low stress conditions, specialized redox-sensing transcription factors (SufR, PerR) rapidly detect and respond to specific ROS signals. Under moderate stress, global regulator RpaB coordinates genome-wide resource reallocation through multiple regulatory states alongside accessory co-regulators. Under severe stress, convergent growth-arrest programs activate with bacterial stringent response signatures.

RpaB emerged as the central coordinator of stress response, with our analysis revealing four independently regulated RpaB-associated iModulons rather than a single regulon. The RpaB∼P iModulon (r = -0.92 with O₂, r = 0.99 with growth) promotes photosynthetic gene expression under favorable conditions; RpaB ROS (r = 0.95 with O₂, r = -0.90 with growth) activates repair and protection systems under stress; RpaABC (r = -0.86 with O₂, r = 0.78 with growth) integrates with circadian regulator RpaA to induce night-like metabolic states during stress; and RpaB ycf46 (r = 0.85 with O₂, r = -0.94 with growth) coordinates severe stress checkpoints. This multi-state architecture, combined with RpaB’s coordination of numerous co-regulators, establishes it as a master molecular switch precisely calibrating growth-survival balance. We next examined how specific iModulons orchestrate responses within each stress regime, beginning with specialized redox sensors mounting initial low-stress defense.

### Redox-Sensing Transcription Factors Drive Initial Stress Defense Through Specialized Regulatory Modules

Low stress conditions (0-140% air saturation) rely on molecular sentinels that detect specific ROS signals and rapidly activate protective systems while maintaining near-normal growth. Among identified metalloregulatory sensors (**Supplementary Table 3**), SufR and PerR emerged as critical <u>first responders</u> with strong correlation across O_2_ levels (r ≥ 0.80) (**Figure 4a**). These transcription factors function as redox-sensitive repressors that de-repress stress-response genes upon ROS sensing[35, 36], enabling rapid antioxidant defense mobilization (**Figure 4b-d**).

**Figure 4.**
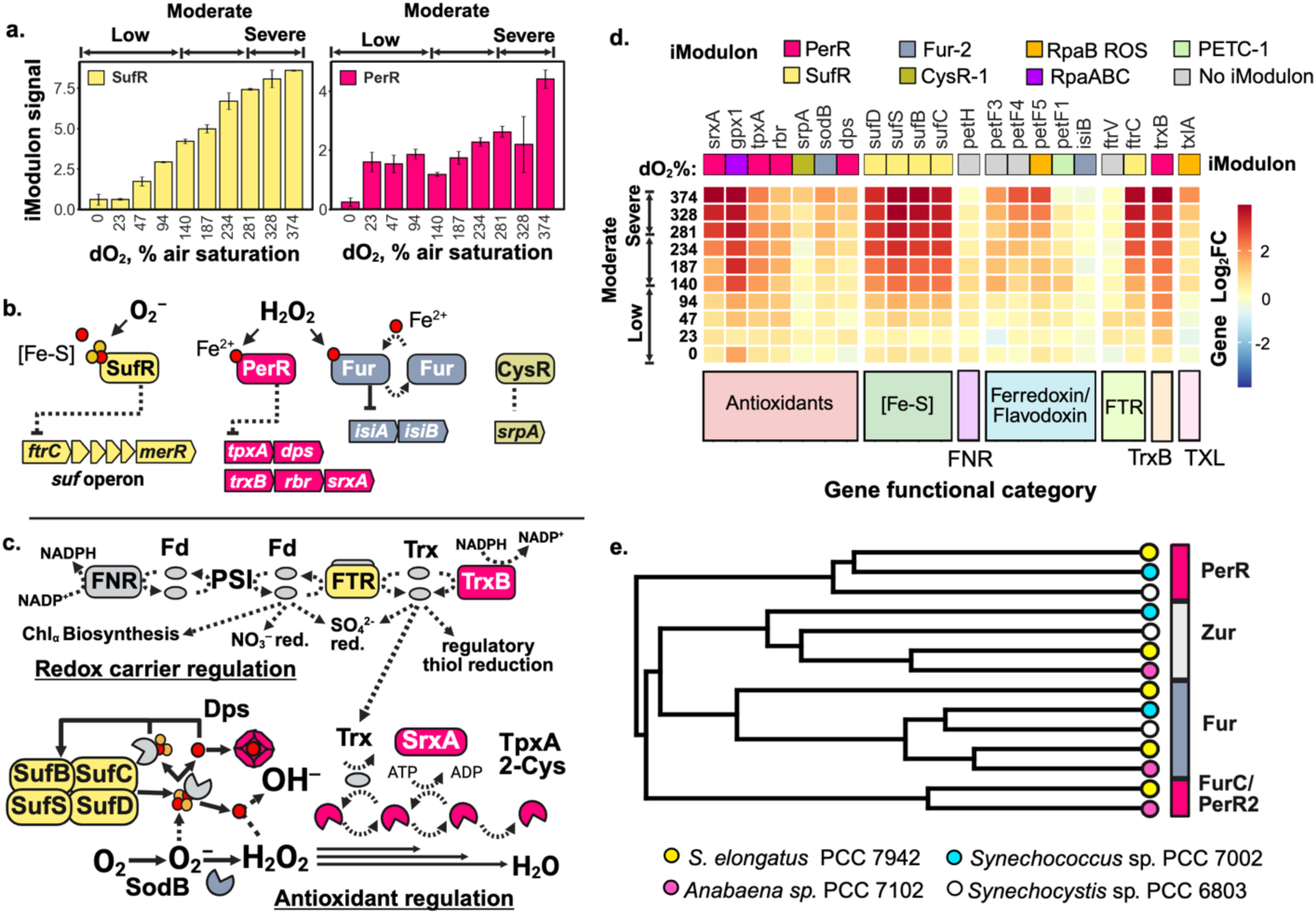
Metalloregulatory sensors SufR and PerR orchestrate primary oxidative stress defense extending through elevated stress phases. **(a)** Dynamic iModulon activity profiles demonstrate immediate SufR activation (r = 0.99) and graduated PerR response (r = 0.80) across the oxygen gradient, revealing hierarchical deployment of redox sensors. **(b)** Molecular mechanisms of redox sensing and transcriptional control: SufR iron-sulfur clusters detect hydroxyl radicals regulating the *suf* operon; PerR iron cofactor senses H₂O₂ controlling thioredoxin reductases, rubrerythrin, and sulfiredoxin; Fur monitors Fe²⁺ availability and H₂O₂ presence governing the *isiAB* iron-stress response. **(c)** Integrated ROS defense network coordinates multiple protective strategies: reducing power diversion to thioredoxin systems (FTR, TrxB), Fe-S cluster maintenance (SufBCDS), iron sequestration preventing Fenton chemistry (Dps), and direct ROS neutralization (SodB, TpxA, Rbr). **(d)** Coordinated expression profiles of redox carriers and antioxidant genes demonstrate multi-layered regulatory integration. Genes are organized by functional categories and labeled with their controlling iModulon, revealing how multiple regulatory modules integrate to mount comprehensive oxidative defense. **(e)** Phylogenetic analysis of Fur family regulators across model cyanobacteria identifies two distinct evolutionary clusters containing PerR iModulon-associated regulators (PerR and PerR2), indicating evolutionary specialization for H₂O₂ sensing in PCC 7942.

The SufR iModulon demonstrated exceptional sensitivity to oxidative stress, having the strongest correlation to oxygen level among all iModulons (r = 0.99). The iModulon signal strongly reflected gene expression activity, explaining 94.7% of variance in the iModulon gene set. This iModulon governs the suf operon (*ftrC-sufBCDS-merR*) with dual protective functions: SufBCDS maintains iron-sulfur cluster integrity, preventing hydroxyl radical formation via Fenton chemistry, while FtrC influences cellular redox balance as a ferredoxin-thioredoxin reductase component [35] (**Figure 4c**). The extraordinary response of the SufR iModulon indicates SufR acts as a highly sensitive redox switch with minimal regulatory crosstalk—ideal for rapid stress detection. SufR activation correlates negatively with growth (r = -0.89), reflecting the metabolic cost of defense.

While SufR responds immediately to oxidative stress, the PerR iModulon shows a more graduated response, with initial activation under low stress conditions (r = 0.40) that intensifies dramatically under moderate stress (r = 0.97). This iModulon controls hydrogen peroxide-specific defenses, explaining 48.1% of its gene set variance. Our analysis identified two distinct PerR homologs forming separate phylogenetic clusters: PerR (Synpcc7942_1648) groups with PCC 6803/7002 regulators[37], while PerR2 (Synpcc7942_1803) clusters with PCC 7120 PerR/FurC[38], suggesting specialized functions (**Figure 4e**).

The PerR regulon comprises a coordinated H₂O₂ defense system including six genes homologous to the PCC 7120 PerR regulon[38] (**Supplementary Table 4**). This system integrates multiple protective mechanisms: ferritin (Dps) for iron sequestration preventing Fenton reactions[39]; H₂O₂-reducing enzymes rubrerythrin (Rbr) and 2-Cys peroxiredoxin (TpxA); sulfiredoxin (SrxA) for reactivating oxidized peroxiredoxins[40]; and NADPH-ferredoxin reductase (TrxB) for channeling reducing power to the thioredoxin system. While both SufR and PerR regulate proteins maintaining thioredoxin activity, the direct link between *trxB* expression and ROS reduction[41] suggests a strategic shift toward enhanced antioxidant defense at moderate stress levels >140% air saturation. Like SufR, PerR activation negatively correlates with growth (r = -0.86), indicating resource allocation toward defense.

This coordinated activation of SufR and PerR creates a sophisticated two-pronged defense strategy. SufR prevents ROS formation by maintaining iron-sulfur clusters and controlling free iron, while PerR manages H₂O₂ detoxification once formed. Both regulators enhance thioredoxin system capacity—SufR through FtrC and PerR through TrxB—creating a unified redox buffer system. This defense architecture is further supported by reprogramming of electron carrier expression under moderate stress: primary carriers (*petH*/FNR and *petF1*) become repressed while alternative ferredoxins (*petF3*, *petF4*, *petF5*) are induced (**Figure 4d**), redirecting reducing power toward antioxidant defense.

Low stress conditions thus represent remarkably effective initial defense—growth continues while robust protective systems activate. However, as stress intensifies, specialized sensors alone cannot maintain homeostasis, necessitating genome-wide resource reallocation from growth to survival through the master regulator RpaB.

### RpaB Master Switch Dynamically Coordinates Growth-Defense Balance Through Multiple Regulatory States

As oxidative stress intensifies (140-280% air saturation), specialized sensors and targeted antioxidant responses cannot maintain homeostasis, requiring genome-wide resource reallocation. RpaB, an OmpR-family response regulator, functions as the central coordinator of this metabolic transition, operating through multiple distinct regulatory states with co-regulator networks across the stress continuum.

RpaB integrates multiple redox signals through sophisticated sensory mechanisms. Histidine kinase NblS phosphorylates RpaB under low-stress conditions and dephosphorylates under high-stress conditions by sensing PSII Q_a_ redox state[13, 20]. Simultaneously, RpaB responds to PSI acceptor side redox state, reflecting intracellular thiol reduction controlled by thioredoxin[42]. ICA analysis revealed how this multi-input sensing enables sophisticated regulatory output: instead of a single regulon, RpaB operates through four independently regulated iModulons, each representing a distinct regulatory state. The first three states—growth-promoting RpaB∼P, stress-protective RpaB ROS, and circadian-integrated RpaABC (co-regulated with RpaA)—show distinct activity patterns and cross-correlations (**Figure 5a-d**), while the severe stress checkpoint RpaB ycf46 is discussed later. Each state exhibits quantifiable correlations with oxygen levels and growth rates, enabling master regulator RpaB to orchestrate stress-level-specific transcriptional cascades through direct control of target genes, five sigma factors, and transcriptional regulators *srrA* and *rpoD*3[43].

**Figure 5.**
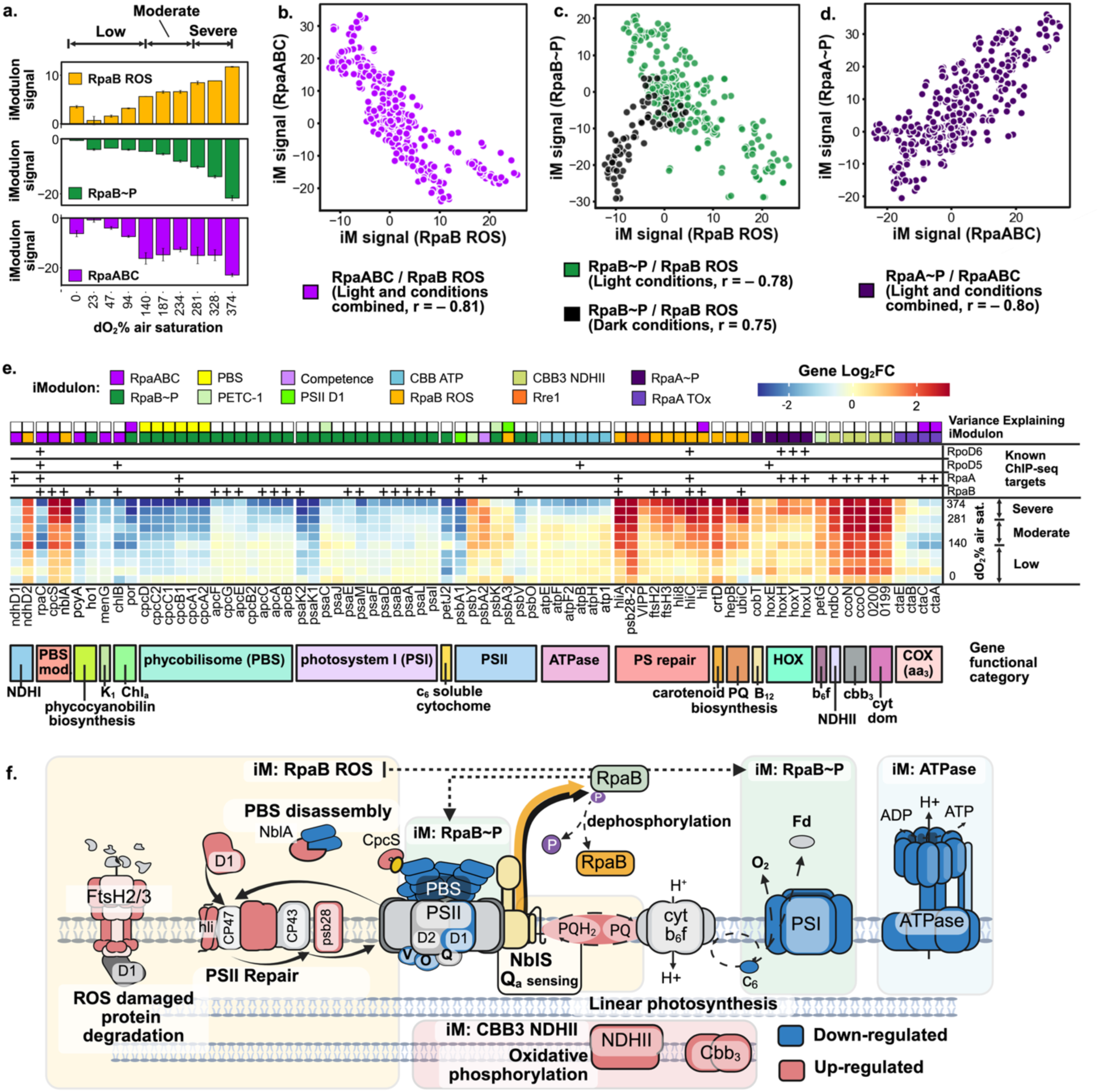
RpaB operates through three independent regulatory states to reprogram photosynthetic electron transport under stress. **(a)** Divergent activity patterns of RpaB-associated iModulons across the oxidative stress gradient reveal state-specific responses. RpaB ROS activates linearly with increasing stress, RpaB∼P shows escalating repression under severe stress, while RpaABC exhibits biphasic repression at moderate and extreme stress levels. **(b-d)** Correlation analyses across 407 RNA-seq samples confirm distinct regulatory state relationships: **(b)** RpaB ROS and RpaABC show inverse coordination (r = -0.81); **(c)** light-dependent switching between RpaB states with RpaB ROS and RpaB∼P showing negative correlation under light (r = - 0.78, green) but positive correlation in darkness (r = 0.75, black); **(d)** strong positive correlation between RpaABC and circadian regulator RpaA∼P (r = 0.80) reveals integration of stress and circadian metabolic programs. **(e)** Comprehensive regulatory map of photosynthetic and electron transport genes showing their variance-explaining iModulons. Analysis includes genes with total variance >0.1 where iModulon(s) explain >30% expression variance, with multiple iModulon assignments revealing complex regulatory integration. **(f)** Model of RpaB-mediated reorganization of photosynthetic and electron transport systems. RpaB ROS activates PSII repair machinery and phycobilisome degradation, while RpaB∼P represses core photosynthetic components. Sophisticated PSII D1 regulation illustrates stress response complexity: growth-associated isoform *psbA1* (PSII D1 iModulon) undergoes continuous suppression with increasing stress, while stress-associated isoforms show divergent regulation—*psbA3* is co-regulated by both RpaB ROS and PSII D1 iModulons, whereas *psbA2* belongs exclusively to the Competence iModulon. This differential D1 isoform regulation enables precise tuning of PSII function across stress levels. Additional adaptations include upregulation of alternative electron transport pathways via type II NDH (NdbC) and *cbb₃* terminal oxidase, providing respiratory bypass routes during photosynthetic stress.

The RpaB ROS iModulon exemplifies this regulatory architecture as the stress defense state (**Figure 5a**), showing strong positive correlation with oxidative stress (r = 0.95) and negative correlation with growth (r = -0.90). This iModulon controls 29 genes (**Supplementary Table 3**), including 17 experimentally confirmed RpaB ChIP-seq targets[43], coordinating a comprehensive stress response system. Key components include PSII repair machinery (*psbA2-3*, *psbD2*, *hli*, *ftsH2/3*) and phycobilisome disassembly (*cpcS*, *nblA*), photoprotective pigment synthesis (*crtD*, *ubiC*), and transcriptional co-regulators *rpoD3* and *srrA* (**Figure 5e, f**). Critical redox regulators: [2Fe-2S] ferredoxin PetF5 and thylakoid thiol:disulfide interchange protein (TxlA)[44] complement this network, directly linking RpaB activity to cellular redox state.

In contrast, the RpaB∼P iModulon promotes photosynthetic growth and shows the highest correlation with growth rate (r = 0.99) among all iModulons (**Figure 3c**). Strongly repressed by oxidative stress (r = -0.92, **Figure 5a**), this iModulon is enriched for photosynthetic proteins (30/38 genes) and includes *rpaB* transcription itself. Under oxidative stress, RpaB∼P repression coordinates downregulation of photosynthetic machinery: PSII extrinsic proteins (*psbO*, *psbV*), soluble cytochrome c6 (*petJ2*), PSI components, and phycobilisome proteins. The PBS phycocyanin operon (Synpcc7942_1048–1053) is simultaneously repressed by a separate regulatory mechanism captured in the PBS iModulon (**Figures 5e, f**). This RpaB∼P-modulated systematic repression of growth-promoting functions directly explains reduced growth observed under moderate stress.

A third regulatory integration—the RpaABC iModulon—connects RpaB signaling with circadian clock programs and shifts cells toward night-like expression during stress (**Figure 4d**). Also repressed by oxidative stress (r = -0.86) and strongly correlated with the primary RpaA∼P iModulon (r = 0.80), this state coordinates expression of sigma factors *sigF1* and *sigF2* (regulate biofilm and competence, respectively[45, 46]), the PBS state-transition regulator *rpaC*[47] and ribosome hibernation factor *hpf*[48]. The RpaABC iModulon also governs biosynthetic pathways for essential electron transport cofactors—chlorophyll, phylloquinol (*menG*), ubiquinone (*coq4*), and lipoate (*lipB*)—suggesting coordinated metabolic downregulation during stress-induced dormancy. Numerous uncharacterized RpaABC targets suggest undiscovered circadian-stress regulatory links.

Analysis across the complete dataset of 407 transcriptome samples revealed dynamic interplay between these regulatory states, providing mechanistic insight into global regulatory patterns governed by RpaB. RpaB ROS and RpaABC signals show strong inverse correlation globally (r = -0.81), validating their negative correlation during oxidative stress (**Figure 5b**). Similarly, RpaB∼P and RpaB ROS show opposing regulatory control under light conditions (r = - 0.78, green) but positive correlation (both repressed) in darkness (r = 0.75, black) (**Figure 5c, Supplementary Figure 6**). Under moderate stress, RpaB mediates switching from photosynthetic growth to repair and protection (**Figures f**), while RpaABC coordinates additional functions particularly when primary states are repressed, including during darkness. This multi-state architecture—rather than simple on-off switching—enables RpaB to fine-tune the growth-defense balance through modulation by the redox-sensing histidine kinase NblS (**Figure 5f**): phosphorylation-dependent RpaB∼P maintains growth programs when possible, dephosphorylated RpaB ROS activates protection when needed, and RpaABC provides regulatory flexibility for circadian and metabolic coordination. This sophisticated mechanism explains how cells maintain reduced but sustainable growth under low and moderate stress, until RpaB ycf46-mediated complete shutdown under severe stress.

### A Network of Local Response Regulators Facilitate Metabolic Rewiring Under Intensifying Stress

While RpaB coordinates cellular responses to moderate stress, an extensive response regulator network systematically reprograms metabolism. Beyond RpaB-mediated photosynthetic control, we identified six response regulator-associated iModulons with significant oxygen correlations: metal homeostasis regulators (ManR, CopR), protein quality control (Rre1), phosphate metabolism (SphR), and circadian regulation (RpaA) (**Table 2**). Each regulator displayed distinctive activation or repression patterns across the stress continuum, most initiating under moderate stress with strongest correlations under severe stress (**Figure 6a-b**). This coordinated response demonstrates how two-component systems integrate redox signals to orchestrate specialized cellular functions essential for stress adaptation.

**Figure 6.**
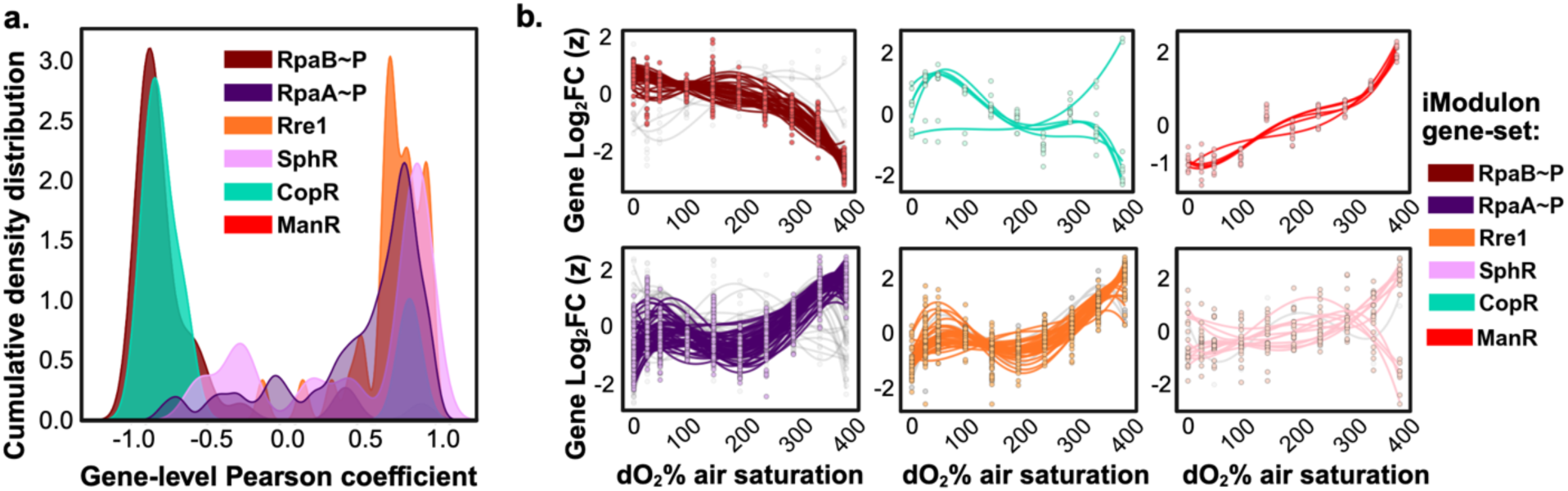
Two-component response regulator-controlled iModulons coordinate specialized functions across the stress gradient, with strongest responses under severe stress. **(a)** Cumulative density distributions of gene-level correlations reveal regulator-specific activation patterns. Analysis includes positively weighted genes within each response regulator iModulon, demonstrating coordinated but specialized regulatory responses. **(b)** Polynomial regression analysis of individual gene expression (z-scores) identifies stress-responsive regulatory targets. Colored genes show strong correlations with oxygen levels above 200% air saturation (|r| > 0.7).

**Table 2.**
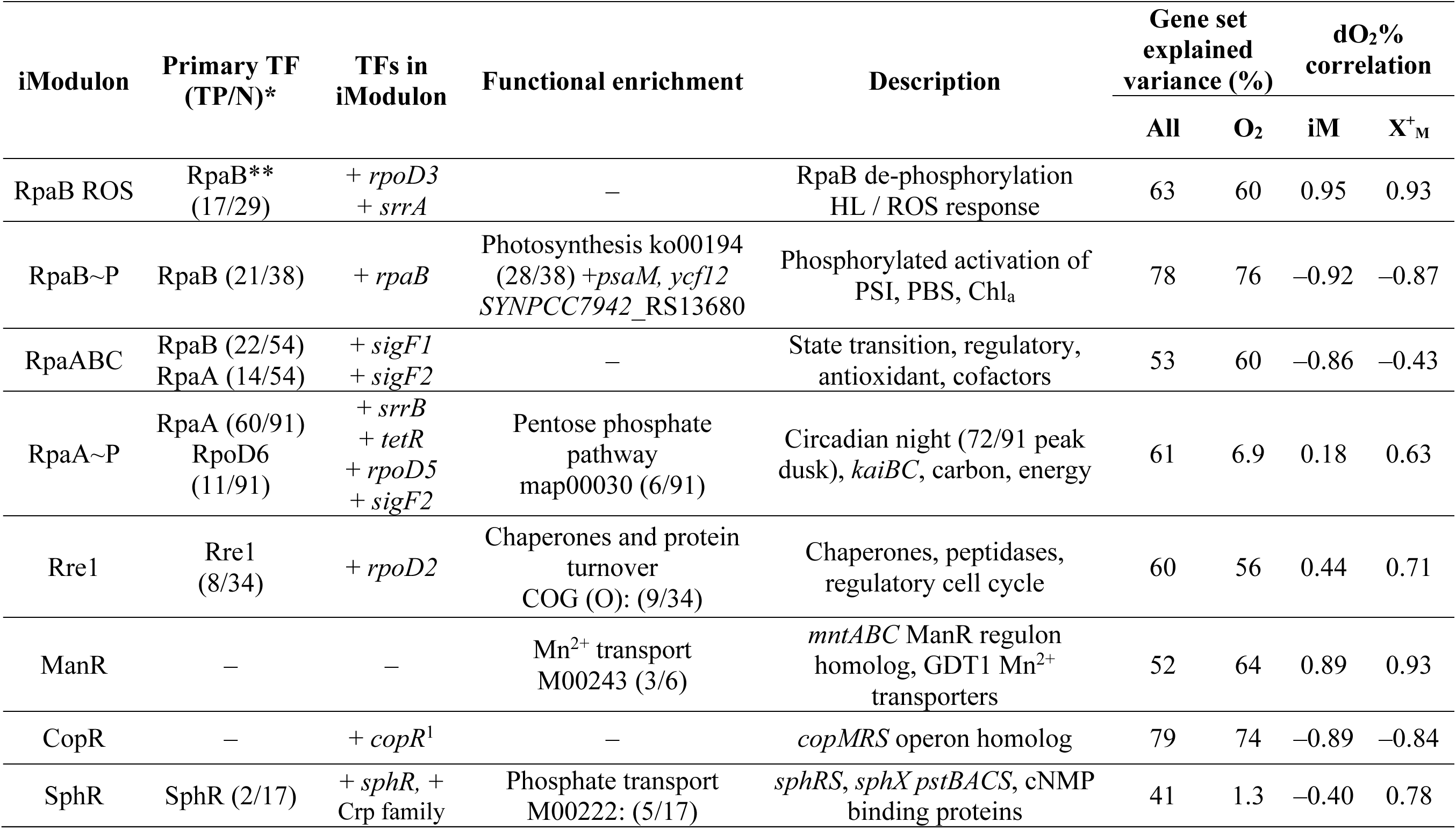
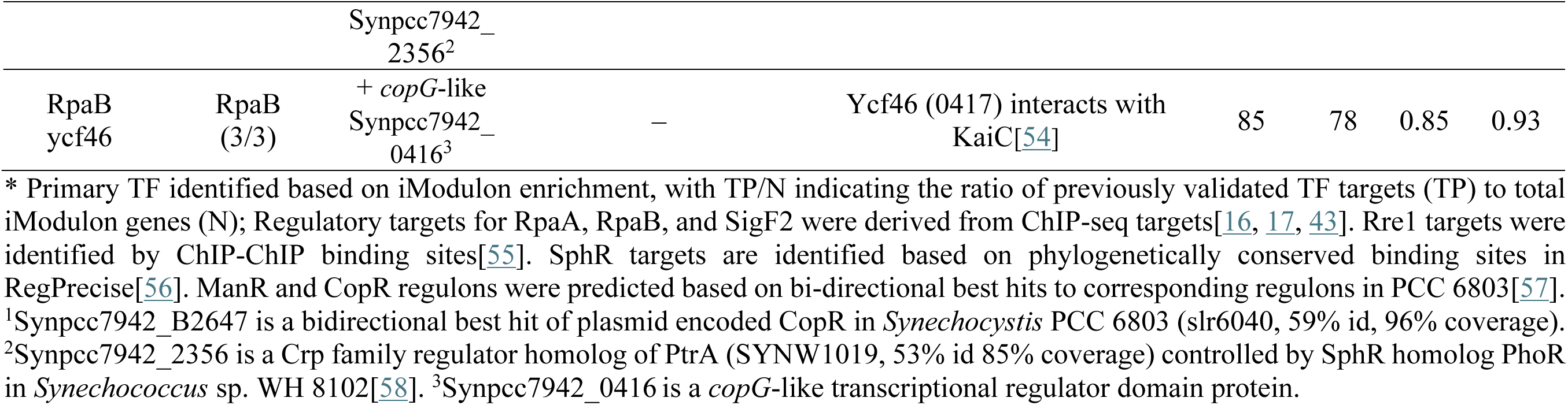
Network of response regulator iModulons coordinates specialized stress adaptations beyond photosynthetic control. Network of response regulator iModulons coordinates specialized stress adaptations beyond photosynthetic control. Six response regulator systems (RpaB, ManR, CopR, Rre1, SphR, RpaA) exhibit distinct activation patterns across the stress continuum. These two-component systems integrate redox signals and control additional transcription and sigma factors within their iModulons to create amplified regulatory cascades controlling metal homeostasis, protein quality maintenance, and carbon metabolism reprogramming. Gene set variance and correlation analyses reveal how this multi-layered regulatory architecture orchestrates specialized cellular functions essential for oxidative stress response.

| iModulon | Primary TF (TP/N)* | TFs in iModulon | Functional enrichment | Description | Gene set explained variance (%) |  | dO<sub>2</sub>% correlation |  |
| --- | --- | --- | --- | --- | --- | --- | --- | --- |
|  |  |  |  |  | All | O<sub>2</sub> | iM | X<sup>+</sup><sub>M</sub> |
| RpaB ROS | RpaB** (17/29) | + *rpoD3*<br>+ *srrA* | — | RpaB de-phosphorylation<br>HL / ROS response | 63 | 60 | 0.95 | 0.93 |
| RpaB~P | RpaB (21/38) | + *rpaB* | Photosynthesis ko00194<br>(28/38) + *psaM*, *ycf12*<br>*SYNPCC7942_RS13680* | Phosphorylated activation of<br>PSI, PBS, Chl<sub>a</sub> | 78 | 76 | −0.92 | −0.87 |
| RpaABC | RpaB (22/54)<br>RpaA (14/54) | + *sigF1*<br>+ *sigF2* | — | State transition, regulatory,<br>antioxidant, cofactors | 53 | 60 | −0.86 | −0.43 |
| RpaA~P | RpaA (60/91)<br>RpoD6 (11/91) | + *srrB*<br>+ *tetR*<br>+ *rpoD5*<br>+ *sigF2* | Pentose phosphate<br>pathway<br>map00030 (6/91) | Circadian night (72/91 peak<br>dusk), *kaiBC*, carbon, energy | 61 | 6.9 | 0.18 | 0.63 |
| Rre1 | Rre1 (8/34) | + *rpoD2* | Chaperones and protein<br>turnover<br>COG (O): (9/34) | Chaperones, peptidases,<br>regulatory cell cycle | 60 | 56 | 0.44 | 0.71 |
| ManR | — | — | Mn<sup>2+</sup> transport<br>M00243 (3/6) | *mntABC* ManR regulon<br>homolog, GDT1 Mn<sup>2+</sup><br>transporters | 52 | 64 | 0.89 | 0.93 |
| CopR | — | + *copR*<sup>1</sup> | — | *copMRS* operon homolog | 79 | 74 | −0.89 | −0.84 |
| SphR | SphR (2/17) | + *sphR*, +<br>Crp family | Phosphate transport<br>M00222: (5/17) | *sphRS*, *sphX* *pstBACS*, cNMP<br>binding proteins | 41 | 1.3 | −0.40 | 0.78 |
|  |  | Synpcc7942_2356<sup>2</sup> |  |  |  |  |  |  |
| RpaB ycf46 | RpaB (3/3) | + *copG*-like Synpcc7942_0416<sup>3</sup> | – | Ycf46 (0417) interacts with KaiC[54] | 85 | 78 | 0.85 | 0.93 |

\* Primary TF identified based on iModulon enrichment, with TP/N indicating the ratio of previously validated TF targets (TP) to total iModulon genes (N); Regulatory targets for RpaA, RpaB, and SigF2 were derived from ChIP-seq targets[16, 17, 43]. Rre1 targets were identified by ChIP-ChIP binding sites[55]. SphR targets are identified based on phylogenetically conserved binding sites in RegPrecise[56]. ManR and CopR regulons were predicted based on bi-directional best hits to corresponding regulons in PCC 6803[57]. <sup>1</sup>Synpcc7942\_B2647 is a bidirectional best hit of plasmid encoded CopR in *Synechocystis* PCC 6803 (slr6040, 59% id, 96% coverage). <sup>2</sup>Synpcc7942\_2356 is a Crp family regulator homolog of PtrA (SYNW1019, 53% id 85% coverage) controlled by SphR homolog PhoR in *Synechococcus* sp. WH 8102[58]. <sup>3</sup>Synpcc7942\_0416 is a *copG*-like transcriptional regulator domain protein.

Among response regulators showing strong oxygen correlations, ManR and CopR emerged as key metal homeostasis coordinators with opposing responses. These iModulons were identified based on homology to well-characterized regulatory systems in PCC 6803[49, 50]. The ManR iModulon displaying strong positive correlation with oxygen levels (r = 0.89) and negative correlation with growth (r = -0.92), controlling expression of manganese transporter homologs *mntCAB* and GDT1-domain manganese transporters. This activation of manganese uptake systems suggests protective strategy, as manganese functions both as a critical cofactor for manganese-dependent superoxide dismutase and as a non-enzymatic antioxidant that can replace iron in metalloproteins to prevent Fenton chemistry-driven hydroxyl radical formation[51].

In contrast, the CopR iModulon showed strong negative correlation with oxygen (r = -0.89) and positive correlation with growth (r = 0.89), indicating repression of the *copMRS* copper response system under oxidative stress. This observation aligns with findings in PCC 6803, where CopR regulates copper homeostasis[50]. The divergent regulation of metal homeostasis systems highlights the need to maintain proper metalloprotein composition during oxidative stress, potentially preventing mismetallation events that could exacerbate ROS damage[51].

The Rre1 iModulon showed significant activation during severe oxidative stress (r = 0.90), with its gene set highly enriched for chaperones and protein turnover functions (COG class O, 9/34 genes). Among regulated genes were essential cell cycle control factors, including cell division inhibitor SulA and Bax inhibitor (Synpcc7942_1820), suggesting coordinated growth arrest during severe stress. Importantly, the Rre1 iModulon signal clustered with PerR in global hierarchical analysis across the entire RNA-seq dataset (**Supplementary Figure 7**), linking activation of both iModulons to the antioxidant defense system and revealing coordinated regulation of protein repair and ROS defense mechanisms.

While some iModulons showed direct oxygen correlations, others revealed complex oxidative stress relationships. The iModulons enriched in RpaA and SphR targets showed relatively poor resolution by ICA signal decomposition under oxidative stress conditions, with the primary RpaA∼P iModulon explaining only 6.9% variance compared to 61% explained variance across all datasets. However, deeper inspection revealed significant gene-level correlations with oxidative stress (**Figure 6a** and **b**). The median gene correlation for positively weighted genes in these iModulon gene sets (X^+^_M_ = 0.63 for RpaA∼P, and 0.78 for SphR) was comparable to well-resolved iModulon gene sets (**Table 4**, **Figure 6a, b**). These iModulons control critical downstream regulators and proteins involved in phosphate assimilation, redox homeostasis, and central carbon metabolism, suggesting their importance in integrated stress responses despite weaker signal resolution.

Collectively, our analysis reveals that oxidative stress drives dynamic genome-wide transcriptional remodeling through coordinated interplay of multiple global and pathway-specific response regulators. These findings are particularly significant given recent evidence that oxidation drives oligomerization of response regulators RpaA, RpaB, and Rre1 in 6803 [52], while thioredoxin reduces disulfide bonds in RpaA, RpaB, and ManR *in vitro*[53]. Such evidence indicates direct mechanistic links between cellular redox state and response regulator activity, revealing opportunities for elucidating redox PTM modulation of two-component systems.

Coordinated ManR and Rre1 activation with CopR repression during severe stress bridges moderate-to-severe growth repression, their iModulons showing strongest growth rate correlations. This multi-regulator response provides independent control over essential pathways, while severe stress triggers ultimate survival mechanisms via complete growth shutdown.

### Severe Oxidative Stress Activates a Stringent Response-Like Checkpoint Mechanism That Systematically Represses Essential Cellular Functions

Severe stress revealed a remarkable finding: oxidative stress activates dual-control mechanisms linking redox sensing to bacterial stringent response. This represents the critical point where defense shifts from active protection to dormancy-like states preserving resources by systematically halting energy-intensive growth. The regulatory architecture governing this transition shows remarkable similarities to the stringent response, a well-characterized bacterial stress adaptation, orchestrated through sophisticated networks involving redox-responsive regulators and iModulons activated or repressed specifically under severe conditions.

Severe stress triggers coordinated repression of multiple iModulons governing fundamental cellular functions essential for growth (**Figure 7a**). This systematic shutdown included: photosynthetic apparatus repression through the PSII D1 iModulon (growth rate correlation, GR-r = 0.97) affecting all PSII D1 isomers (*psbA1-3*) and the ATPase iModulon (GR-r = 0.87) repressing F₀F₁ ATP synthase; carbon fixation machinery shutdown via the CCM-1 iModulon (GR-r = 0.96) repressing carboxysome and RuBisco operons (*ccmK2LMN*, *rbsLS*) critical for photosynthetic carbon assimilation; protein synthesis cessation through two ribosomal iModulons—Ribo-1 (GR-r = 0.93) and Ribo-2 (GR-r = 0.91)—causing widespread ribosomal protein repression and effectively halting cellular protein synthesis capacity; and cell division arrest via the Cell Div iModulon (GR-r = 0.93) repressing cell division genes *cdv2* and *spollD*. This coordinated repression of multiple growth-essential cellular systems demonstrates comprehensive physiological shutdown in response to severe oxidative stress that cannot be addressed through defensive mechanisms activated in earlier stress conditions. The scale of this transcriptional reprogramming was evident in the dramatic increase in differentially expressed genes under severe stress compared to earlier regimes (**Supplementary Figure 3**).

**Figure 7.**
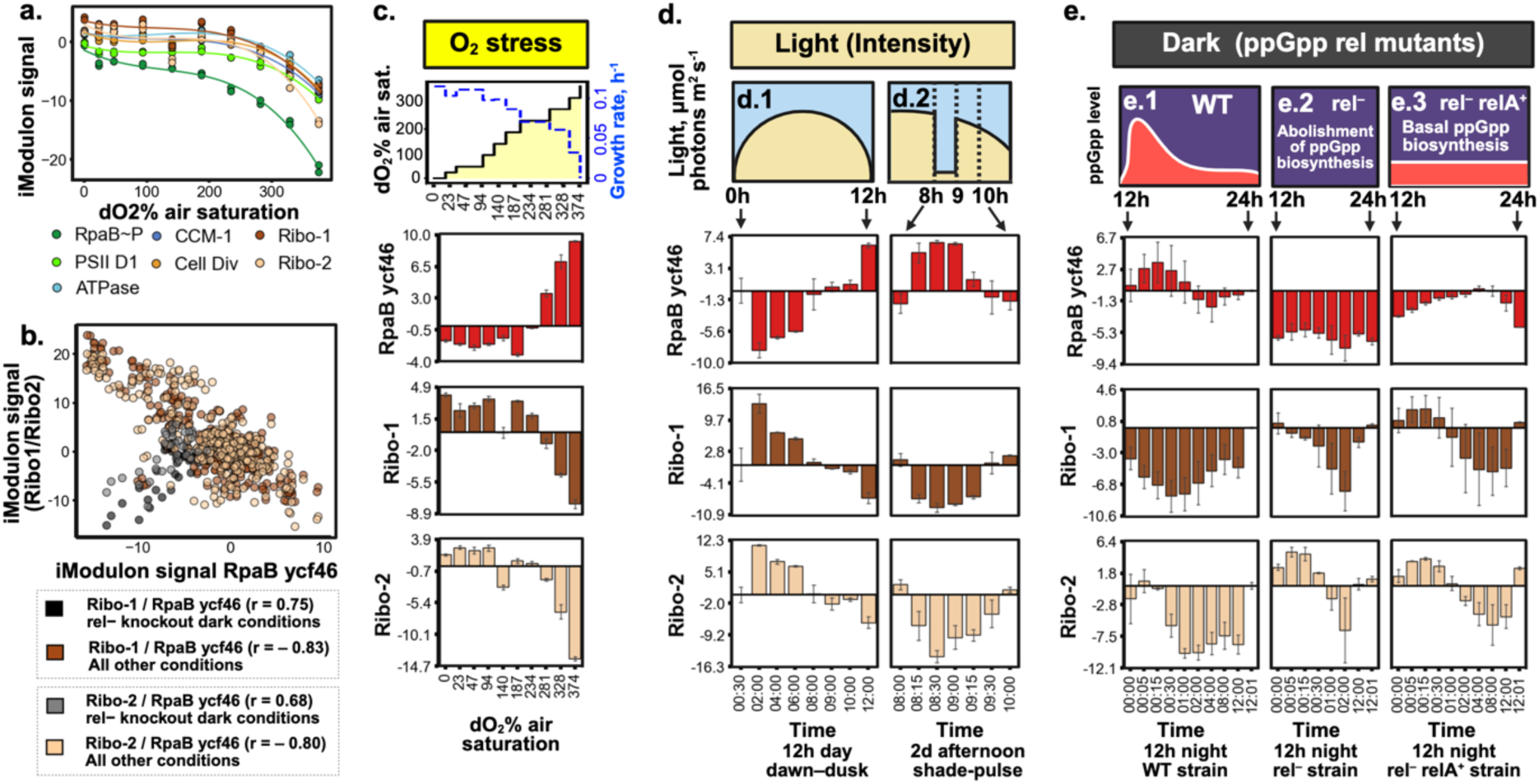
Severe oxidative stress activates stringent response-dependent arrest of critical cellular functions through the RpaB ycf46 iModulon checkpoint. **(a)** Growth-essential iModulons exhibit systematic repression during severe oxidative stress correlating with growth inhibition and demonstrating coordinated shutdown of core cellular processes: photosynthesis (PSII D1), carbon fixation (CCM-1), protein synthesis (Ribo-1, Ribo-2), and cell division (Cell Div). **(b)** RpaB ycf46 iModulon exhibits strict anti-correlation with ribosomal iModulons (Ribo-1 and Ribo-2) across all conditions except (p)ppGpp synthase mutants in darkness (rel^−^ and rel^−^ + relA^E335Q^; from Puszynska et al., 2017)[48]. **(c)** Severe stress triggers reciprocal regulation: sharp RpaB ycf46 activation coincides with robust ribosomal gene repression. **(d)** Light reduction triggers RpaB ycf46 checkpoint during **(d.1)** natural dusk transitions and **(d.2)** acute experimental shading, revealing shared stringent response pathways between severe oxidative stress and acute light limitation. **(e)** Time-course analysis demonstrates stringent response requirement: **(e.1)** wild-type cells show transient RpaB ycf46 activation at light-dark transition that subsides over 12 hours; **(e.2)** complete loss of RpaB ycf46 activity and altered ribosome dynamics in ppGpp-deficient rel⁻ mutant; (**e.3**) partial restoration with basal (p)ppGpp in rel⁻ relA⁺ strain confirms dose-dependent regulation.

Beyond this systematic repression, severe stress is marked by strong induction of three key iModulons showing the highest correlations with oxygen stress (r > 0.96) and specifically linked to growth suppression: RpaB ycf46 (GR-r = -0.94), the largely uncharacterized sigG rpoD2 (GR-r = -0.99), and pyrR dnaQ (r = 0.96). These severe stress-specific iModulons function in concert with the redox-responsive regulators described previously—including ManR, Rre1, and CopR— which acquire critical growth control functions. Specifically, the CopR iModulon signal clusters with the sigG rpoD2 signal in global hierarchical analysis across the entire RNA-seq dataset (**Supplementary Figure 7**), indicating convergent regulatory control during the transition to severe stress conditions.

Among these stress-activated iModulons, RpaB ycf46 emerged as a critical regulatory hub linking oxidative stress perception to cellular growth arrest. Despite controlling a single three-gene operon (Synpcc7942_0416-0418), this iModulon explains 1.3% of global gene expression variance, ranking it among the most influential regulatory signals in our dataset (**Supplementary Data 1**). The operon encodes a KaiC-interacting AAA family ATPase (Ycf46, Synpcc7942_0417[54]) previously shown to affect circadian timing in PCC 7942 and carbon concentration mechanisms in PCC 6803[59].

The RpaB ycf46 iModulon consistently shows strong inverse correlation with ribosomal protein expression across diverse conditions (**Figure 7c-e**). When RpaB ycf46 is activated— during severe oxidative stress, light reduction, or nighttime—ribosomal gene expression is simultaneously repressed, indicating coordinated shutdown of protein synthesis machinery. This anti-correlation pattern suggests that the ycf46 operon functions as a molecular switch that signals the cell to halt growth-related processes.

To understand the mechanism behind this growth control, we examined data from previous studies using mutant strains defective in stringent response regulation[48]. The stringent response is a well-characterized bacterial stress adaptation where cells produce an alarm signal molecule guanosine tetraphosphate/pentaphosphate or (p)ppGpp, that globally represses growth-related genes while activating stress survival genes.

Our ICA analysis demonstrated interesting dependency: in strains lacking the enzyme producing (p)ppGpp (rel⁻ knockout strains), the RpaB ycf46 iModulon is completely repressed, even under conditions that normally activate it (**Figure 7d**, **Figure 7e.2**). This demonstrates that (p)ppGpp is essential for activation of the RpaB ycf46 iModulon. Furthermore, when low levels of (p)ppGpp production are restored (rel⁻ relA⁺ strains), the RpaB ycf46’s normal regulatory pattern returns, but full activation still requires the elevated (p)ppGpp levels (**Figure 7e.3**), that occurre during stress conditions.

These findings indicate that the RpaB ycf46 iModulon operates under dual regulatory control: it requires both RpaB activation (responding to redox stress) and elevated (p)ppGpp levels (the stringent response signal). This dual dependency creates a sophisticated checkpoint system where severe oxidative stress simultaneously triggers RpaB-mediated redox sensing and (p)ppGpp-mediated stringent response, converging on the ycf46 operon to coordinate growth shutdown. This stringent response connection reveals unified regulatory architecture linking oxidative stress adaptation to fundamental bacterial stress biology.

### A Unified Multi-Phase Regulatory Model: From ROS Sensing and Metabolic Rebalancing to Global Survival Programs

Integrating our discoveries reveals a quantitative systems-level regulatory framework advancing understanding of how photosynthetic organisms balance growth and survival under oxidative stress **(Figure 8)**.

**Figure 8.**
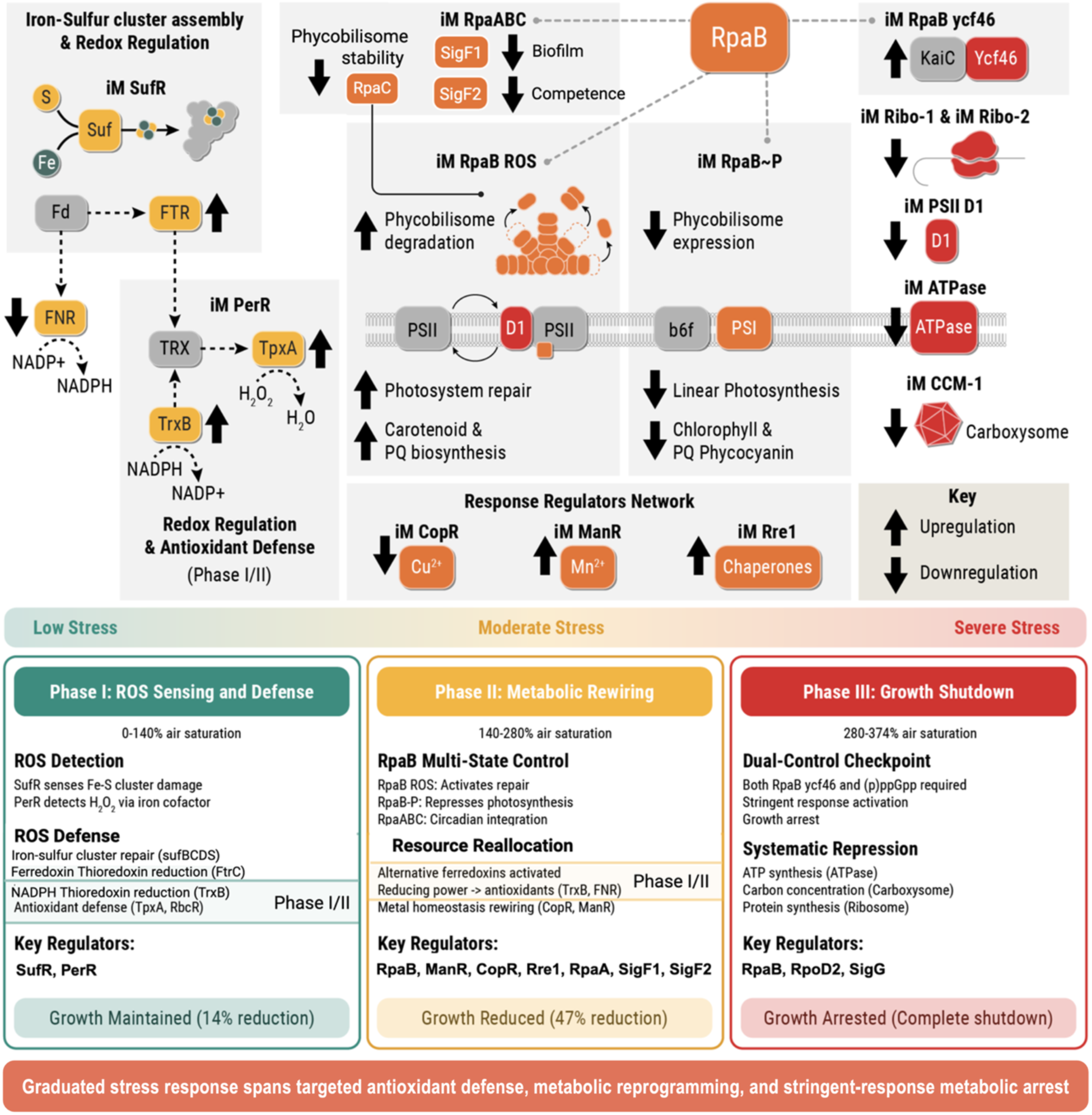
Conceptual three-phase model describes how multi-layered regulatory architecture enables *S. elongatus* PCC 7942 to coordinate dynamic response across oxidative stress continuum through integrated ROS sensors, multi-state global and local response regulators, and stringent response pathways for growth-survival balance. Three regulatory phases provide a simplified framework capturing key transitions within the continuous stress response spectrum across escalating oxidative stress. **Phase I with low stress level (0-140% O₂ saturation):** Specialized metalloregulatory sensors (SufR, PerR) deploy targeted antioxidant defenses while maintaining 86% baseline growth through specific ROS signal detection. **Phase II with moderate stress level (140-280% O₂):** Master regulator RpaB orchestrates genome-wide metabolic reprogramming via three independent states (RpaB∼P, RpaB ROS, RpaABC), systematically reallocating resources from photosynthesis to protection systems. **Phase III with severe stress level (280-374% O₂):** Dual-control checkpoint, requiring both RpaB ycf46 activation and stringent response (p)ppGpp signaling, triggers coordinated shutdown of photosynthesis, carbon fixation, translation, and cell division. This graduated regulatory architecture enables precise calibration of growth-survival trade-offs across the complete stress continuum, revealing integration of photosynthetic redox regulation with fundamental bacterial stress response pathways.

#### Low stress activates specialized ROS sensing and defense systems

Metalloregulatory sensors SufR and PerR function as molecular switches detecting specific ROS signals. SufR responds to Fe-S cluster damage by activating iron sequestration and cluster repair systems, while PerR senses H₂O₂ through its iron cofactor, coordinating ferritin, peroxiredoxins, and thioredoxin reductase expression for comprehensive peroxide defense. This enables near-normal growth while establishing robust antioxidant defenses.

#### Moderate stress activates RpaB-mediated metabolic rewiring

RpaB coordinates resource reallocation through four distinct regulatory states: RpaB ROS (activates repair/protection), RpaB∼P (represses photosynthesis), RpaABC (coordinates with circadian regulator RpaA), and RpaB ycf46 (coordinates severe stress checkpoints). This multi-state regulation creates sophisticated switches redirecting resources from growth-oriented photosynthesis toward protective functions. Cells simultaneously replace primary ferredoxins with alternative isoforms, redirecting reducing power toward antioxidant systems. Beyond RpaB, a network of response regulators (ManR, CopR, Rre1) coordinates metal homeostasis, protein quality control, and growth regulation.

#### Severe stress reveals additional key finding: dual-control growth shutdown

Severe stress activates a checkpoint integrating redox sensing with stringent response regulation. The RpaB ycf46 operon requires dual activation—RpaB-mediated redox sensing plus elevated (p)ppGpp levels. Evidence from rel⁻ mutants demonstrates complete ycf46 abolition without (p)ppGpp, confirming this dual dependency. When activated, ycf46 coordinates systematic repression of photosynthesis, carbon fixation, protein synthesis, and energy generation while multiple response regulators simultaneously coordinate specialized protective functions.

This graduated regulatory architecture represents a multi-scale network solution to oxygenic photosynthesis challenges. Low stress establishes ROS detoxification; moderate stress adds resource reallocation through RpaB’s multiple states and coordinated response regulator networks; severe stress introduces stringent response checkpoints ensuring survival under overwhelming stress. The system shows striking parallels with circadian regulation—RpaABC coordinates night-like expression states during stress, demonstrating how redox signals interface with temporal regulation.

This work demonstrates how integrative systems biology approaches, combining controlled perturbation experiments with comprehensive transcriptomic network analysis, reveal quantitative regulatory principles governing cellular adaptation. The graduated response ensures optimal resource utilization: maintaining growth when manageable, reallocating when adaptable, prioritizing survival when overwhelming. Beyond fundamental advances, these discoveries address critical biotechnology challenges where stress tolerance limits photosynthesis-based bioprocess efficiency. The modular network architecture identified here—from specialized sensors through multi-state coordinators to stringent checkpoints—provides systematic engineering strategies for rational biodesign of robust cyanobacterial platforms. This regulatory framework offers precision tools for synthetic biology applications and predictive capabilities for cyanobacterial responses to increasing environmental stressors, transforming understanding of photosynthetic survival in our oxygen-rich world.

## Acknowledgements

We thank Dr. Austin Gluth for biological insights and helpful discussions, and Dr. Bin Yang for editorial comments. Figures in this manuscript were created using Biorender. The research described in this paper is part of the Predictive Phenomics Initiative and was conducted under the Laboratory Directed Research and Development Program at Pacific Northwest National Laboratory. This work is also partially supported by the NW-BRaVE for Biopreparedness project funded by the U. S. Department of Energy (DOE), Office of Science, Office of Biological and Environmental Research, under FWP 81832. A portion of this research was performed on a project award (https://doi.org/10.46936/staf.proj.2023.61054/60012367) from the Environmental Molecular Sciences Laboratory, a DOE Office of Science User Facility sponsored by the Biological and Environmental Research program (contract No. DE-AC05-76RL01830). Pacific Northwest National Laboratory is a multi-program national laboratory operated by Battelle for the DOE under Contract DE-AC05-76RLO 1830.

## Author contributions

P.B. and M.S.C. funding acquisition and project administration; P.B. conception and design; E.H. photobioreactor setup and maintenance; P.B. oxidative stress experiments; N.C.S. morphological analysis through confocal microscopy; P.B. processing and interpretation of cultivation data; M.G. RNA extraction and manipulation; A.G. processing and preliminary analysis of transcriptome data; Z.J. iModulon analysis; Z.J. and P.B. data visualization and interpretation; Z.J. and P.B. writing – original draft; P.B., Z.J., W.-J.Q., and M.S.C. writing – review & editing.

## Authors agreement to authorship and submission

All persons designated as author agree to submit the manuscript for peer review.

## Code Availability

GitHub link

## Data availability

Primary RNA-Seq raw measurement data are openly accessible for download at the Gene Expression Omnibus (GEO) community repository under the accession GSE288532, GSE288532. Processed transcriptomics datasets, containing normalized counts and differential gene expression results files and experimental design metadata, are openly accessible for download at the PNNL DataHub Predictive Phenomics Initiative (PPI) Project dataset catalog under https://doi.org/10.25584/2510500.

## Statement of informed consent, human/animal rights

No conflicts, informed consent, human or animal rights applicable.

## References

1. Davies, K.J.A., Oxidative stress: the paradox of aerobic life. Biochemical Society Symposia, 1995. 61: p. 1–31.

2. Imlay, J.A., Pathways of oxidative damage. Annual Reviews in Microbiology, 2003. 57(1): p. 395–418.

3. Liu, X., J. Sheng, and R. Curtiss Iii, Fatty acid production in genetically modified cyanobacteria. Proceedings of the National Academy of Sciences, 2011. 108(17): p. 6899–6904.

4. Lan, E.I. and J.C. Liao, ATP drives direct photosynthetic production of 1-butanol in cyanobacteria. Proceedings of the National Academy of Sciences, 2012. 109(16): p. 6018–6023.

5. Pospíšil, P., Production of Reactive Oxygen Species by Photosystem II as a Response to Light and Temperature Stress. Frontiers in Plant Science, 2016. 7.

6. Hamilton, T.L., The trouble with oxygen: The ecophysiology of extant phototrophs and implications for the evolution of oxygenic photosynthesis. Free Radical Biology and Medicine, 2019. 140: p. 233–249.

7. Zhang, N., E.M. Mattoon, W. McHargue, B. Venn, D. Zimmer, K. Pecani, J. Jeong, C.M. Anderson, C. Chen, J.C. Berry, M. Xia, S.-C. Tzeng, E. Becker, L. Pazouki, B. Evans, F. Cross, J. Cheng, K.J. Czymmek, M. Schroda, T. Mühlhaus, and R. Zhang, Systems-wide analysis revealed shared and unique responses to moderate and acute high temperatures in the green alga Chlamydomonas reinhardtii. Communications Biology, 2022. 5(1): p. 460.

8. Masuda, K., T. Sakurai, and A. Hirano, A coupled model between circadian, cell-cycle, and redox rhythms reveals their regulation of oxidative stress. Scientific Reports, 2024. 14(1): p. 15479.

9. Diamond, S., B.E. Rubin, R.K. Shultzaberger, Y. Chen, C.D. Barber, and S.S. Golden, Redox crisis underlies conditional light–dark lethality in cyanobacterial mutants that lack the circadian regulator, RpaA. Proceedings of the National Academy of Sciences, 2017. 114(4): p. E580–E589.

10. Venkiteswaran, J.J., L.I. Wassenaar, and S.L. Schiff, Dynamics of dissolved oxygen isotopic ratios: a transient model to quantify primary production, community respiration, and air–water exchange in aquatic ecosystems. Oecologia, 2007. 153(2): p. 385–398.

11. Schmitt, F.-J., G. Renger, T. Friedrich, V.D. Kreslavski, S.K. Zharmukhamedov, D.A. Los, V.V. Kuznetsov, and S.I. Allakhverdiev, Reactive oxygen species: Re-evaluation of generation, monitoring and role in stress-signaling in phototrophic organisms. Biochimica et Biophysica Acta (BBA) - Bioenergetics, 2014. 1837(6): p. 835–848.

12. Zavřel, T., M. Faizi, C. Loureiro, G. Poschmann, K. Stühler, M. Sinetova, A. Zorina, R. Steuer, and J. Červený, Quantitative insights into the cyanobacterial cell economy. eLife, 2019. 8: p. e42508.

13. Hanaoka, M. and K. Tanaka, Dynamics of RpaB–promoter interaction during high light stress, revealed by chromatin immunoprecipitation (ChIP) analysis in Synechococcus elongatus PCC 7942. The Plant Journal, 2008. 56(2): p. 327–335.

14. Hanaoka, M., N. Takai, N. Hosokawa, M. Fujiwara, Y. Akimoto, N. Kobori, H. Iwasaki, T. Kondo, and K. Tanaka, RpaB, another response regulator operating circadian clock-dependent transcriptional regulation in Synechococcus elongatus PCC 7942. J Biol Chem, 2012. 287(31): p. 26321–7.

15. Gutu, A. and Erin K. O’Shea, Two Antagonistic Clock-Regulated Histidine Kinases Time the Activation of Circadian Gene Expression. Molecular Cell, 2013. 50(2): p. 288–294.

16. Markson, S., Joseph, R. Piechura, Joseph, M. Puszynska, Anna, and K. O’Shea, Erin, Circadian Control of Global Gene Expression by the Cyanobacterial Master Regulator RpaA. Cell, 2013. 155(6): p. 1396–1408.

17. Fleming, K., A Clock-Phased Sigma Factor Cascade Is Required for Global Circadian Transcriptional Rhythms in Cyanobacteria, in The Department of Molecular and Cellular Biology. 2017, Harvard University.

18. Yuan, Y., T. Al Bulushi, A.V. Sastry, C. Sancar, R. Szubin, S.S. Golden, and B.O. Palsson, Machine learning reveals the transcriptional regulatory network and circadian dynamics of Synechococcus elongatus PCC 7942. Proceedings of the National Academy of Sciences, 2024. 121(38): p. e2410492121.

19. Johnson, Z., D. Anderson, M.S. Cheung, and P. Bohutskyi, Gene network centrality analysis identifies key regulators coordinating day-night metabolic transitions in Synechococcus elongatus PCC 7942 despite limited accuracy in predicting direct regulator-gene interactions. Frontiers in Microbiology, 2025. **Volume** 16 **-** 2025.

20. Tsurumaki, T. and K. Tanaka, An Evolutionary Conserved Multi-Stress Sensory Histidine Kinase NblS Associates With Photosystem II Proteins And Responds To Its Redox Status In The Cyanobacterium Synechococcus elongatus PCC 7942. bioRxiv, 2025: p. 2025.01.21.633742.

21. Bairagi, N., S. Watanabe, K. Nimura-Matsune, K. Tanaka, T. Tsurumaki, S. Nakanishi, and K. Tanaka, Conserved Two-component Hik2–Rre1 Signaling Is Activated Under Temperature Upshift and Plastoquinone-reducing Conditions in the Cyanobacterium Synechococcus elongatus PCC 7942. Plant and Cell Physiology, 2022. 63(2): p. 176–188.

22. Saelens, W., R. Cannoodt, and Y. Saeys, A comprehensive evaluation of module detection methods for gene expression data. Nature Communications, 2018. 9(1): p. 1090.

23. Mcconn, J.L., C.R. Lamoureux, S. Poudel, B.O. Palsson, and A.V. Sastry, Optimal dimensionality selection for independent component analysis of transcriptomic data. BMC Bioinformatics, 2021. 22(1).

24. Abramson, B.W., B. Kachel, D.M. Kramer, and D.C. Ducat, Increased Photochemical Efficiency in Cyanobacteria via an Engineered Sucrose Sink. Plant and Cell Physiology, 2016. 57(12): p. 2451–2460.

25. Santos-Merino, M., A. Torrado, G.A. Davis, A. Röttig, T.S. Bibby, D.M. Kramer, and D.C. Ducat, Improved photosynthetic capacity and photosystem I oxidation via heterologous metabolism engineering in cyanobacteria. Proceedings of the National Academy of Sciences, 2021. 118(11).

26. Ducat, D.C., J.A. Avelar-Rivas, J.C. Way, and P.A. Silver, Rerouting Carbon Flux To Enhance Photosynthetic Productivity. Applied and Environmental Microbiology, 2012. 78(8): p. 2660–2668.

27. Melnicki, M.R., G.E. Pinchuk, E.A. Hill, L.A. Kucek, S.M. Stolyar, J.K. Fredrickson, A.E. Konopka, and A.S. Beliaev, Feedback-controlled LED photobioreactor for photophysiological studies of cyanobacteria. Bioresource Technology, 2013. 134: p. 127–133.

28. Liao, Y., G.K. Smyth, and W. Shi, The R package Rsubread is easier, faster, cheaper and better for alignment and quantification of RNA sequencing reads. Nucleic Acids Research, 2019. 47(8): p. e47–e47.

29. O’Leary, N.A., M.W. Wright, J.R. Brister, S. Ciufo, D. Haddad, R. McVeigh, B. Rajput, B. Robbertse, B. Smith-White, D. Ako-Adjei, A. Astashyn, A. Badretdin, Y. Bao, O. Blinkova, V. Brover, V. Chetvernin, J. Choi, E. Cox, O. Ermolaeva, C.M. Farrell, T. Goldfarb, T. Gupta, D. Haft, E. Hatcher, W. Hlavina, V.S. Joardar, V.K. Kodali, W. Li, D. Maglott, P. Masterson, K.M. McGarvey, M.R. Murphy, K. O’Neill, S. Pujar, S.H. Rangwala, D. Rausch, L.D. Riddick, C. Schoch, A. Shkeda, S.S. Storz, H. Sun, F. Thibaud-Nissen, I. Tolstoy, R.E. Tully, A.R. Vatsan, C. Wallin, D. Webb, W. Wu, M.J. Landrum, A. Kimchi, T. Tatusova, M. DiCuccio, P. Kitts, T.D. Murphy, and K.D. Pruitt, Reference sequence (RefSeq) database at NCBI: current status, taxonomic expansion, and functional annotation. Nucleic Acids Res, 2016. 44(D1): p. D733–45.

30. Vijayan, V., I.H. Jain, and E.K. O’Shea, A high resolution map of a cyanobacterial transcriptome. Genome biology, 2011. 12(5): p. R47.

31. Sastry, A.V., Y. Gao, R. Szubin, Y. Hefner, S. Xu, D. Kim, K.S. Choudhary, L. Yang, Z.A. King, and B.O. Palsson, The Escherichia coli transcriptome mostly consists of independently regulated modules. Nature Communications, 2019. 10(1): p. 5536.

32. Zwietering, M.H., I. Jongenburger, F.M. Rombouts, and K. van ‘t Riet, Modeling of the Bacterial Growth Curve. Applied and Environmental Microbiology, 1990. 56(6): p. 1875–1881.

33. Levenspiel, O., The Monod equation: A revisit and a generalization to product inhibition situations. Biotechnology and Bioengineering, 2004. 22(8): p. 1671–1687.

34. Goutelle, S., M. Maurin, F. Rougier, X. Barbaut, L. Bourguignon, M. Ducher, and P. Maire, The Hill equation: a review of its capabilities in pharmacological modelling. Fundamental & Clinical Pharmacology, 2008. 22(6): p. 633–648.

35. Nodop, A., D. Pietsch, R. Höcker, A. Becker, E.K. Pistorius, K. Forchhammer, and K.P. Michel, Transcript profiling reveals new insights into the acclimation of the mesophilic fresh-water cyanobacterium Synechococcus elongatus PCC 7942 to iron starvation. Plant Physiol, 2008. 147(2): p. 747–63.

36. Shen, G., R. Balasubramanian, T. Wang, Y. Wu, L.M. Hoffart, C. Krebs, D.A. Bryant, and J.H. Golbeck, SufR Coordinates Two [4Fe-4S]2+, 1+ Clusters and Functions as a Transcriptional Repressor of the sufBCDS Operon and an Autoregulator of sufR in Cyanobacteria*. Journal of Biological Chemistry, 2007. 282(44): p. 31909–31919.

37. Li, H., A.K. Singh, L.M. McIntyre, and L.A. Sherman, Differential gene expression in response to hydrogen peroxide and the putative PerR regulon of Synechocystis sp. strain PCC 6803. Journal of bacteriology, 2004. 186(11): p. 3331–3345.

38. Sarasa-Buisan, C., J. Guío, M.L. Peleato, M.F. Fillat, and E. Sevilla, Expanding the FurC (PerR) regulon in Anabaena (Nostoc) sp. PCC 7120: Genome-wide identification of novel direct targets uncovers FurC participation in central carbon metabolism regulation. PLOS ONE, 2023. 18(8): p. e0289761.

39. Howe, C., V.K. Moparthi, F.M. Ho, K. Persson, and K. Stensjö, The Dps4 from Nostoc punctiforme ATCC 29133 is a member of His-type FOC containing Dps protein class that can be broadly found among cyanobacteria. PLoS One, 2019. 14(8): p. e0218300.

40. Boileau, C., L. Eme, C. Brochier-Armanet, A. Janicki, C.-C. Zhang, and A. Latifi, A eukaryotic-like sulfiredoxin involved in oxidative stress responses and in the reduction of the sulfinic form of 2-Cys peroxiredoxin in the cyanobacterium Anabaena PCC 7120. New Phytologist, 2011. 191(4): p. 1108–1118.

41. Hishiya, S., W. Hatakeyama, Y. Mizota, N. Hosoya-Matsuda, K. Motohashi, M. Ikeuchi, and T. Hisabori, Binary Reducing Equivalent Pathways Using NADPH-Thioredoxin Reductase and Ferredoxin-Thioredoxin Reductase in the Cyanobacterium Synechocystis sp. Strain PCC 6803. Plant and Cell Physiology, 2008. 49(1): p. 11–18.

42. Kato, N., K. Iwata, T. Kadowaki, K. Sonoike, and Y. Hihara, Dual Redox Regulation of the DNA-Binding Activity of the Response Regulator RpaB in the Cyanobacterium Synechocystis sp. PCC 6803. Plant and Cell Physiology, 2022. 63(8): p. 1078–1090.

43. Piechura, J.R., K. Amarnath, and E.K. O’Shea, Natural changes in light interact with circadian regulation at promoters to control gene expression in cyanobacteria. eLife, 2017. 6.

44. Collier, J.L. and A.R. Grossman, Disruption of a gene encoding a novel thioredoxin-like protein alters the cyanobacterial photosynthetic apparatus. Journal of Bacteriology, 1995. 177(11): p. 3269–3276.

45. Taton, A., C. Erikson, Y. Yang, B.E. Rubin, S.A. Rifkin, J.W. Golden, and S.S. Golden, The circadian clock and darkness control natural competence in cyanobacteria. Nature Communications, 2020. 11(1): p. 1688.

46. Simkovsky, R., R. Parnasa, J. Wang, E. Nagar, E. Zecharia, S. Suban, Y. Yegorov, B. Veltman, E. Sendersky, R. Schwarz, and S.S. Golden, Transcriptomic and Phenomic Investigations Reveal Elements in Biofilm Repression and Formation in the Cyanobacterium Synechococcus elongatus PCC 7942. Frontiers in Microbiology, 2022. 13.

47. Joshua, S. and C.W. Mullineaux, The rpaC gene product regulates phycobilisome-photosystem II interaction in cyanobacteria. Biochim Biophys Acta, 2005. 1709(1): p. 58–68.

48. Puszynska, A.M. and E.K. O’Shea, ppGpp Controls Global Gene Expression in Light and in Darkness in S. elongatus. Cell Reports, 2017. 21(11): p. 3155–3165.

49. Yamaguchi, K., I. Suzuki, H. Yamamoto, A. Lyukevich, I. Bodrova, D.A. Los, I. Piven, V. Zinchenko, M. Kanehisa, and N. Murata, A Two-Component Mn2+-Sensing System Negatively Regulates Expression of the mntCAB Operon in Synechocystis. The Plant Cell, 2002. 14(11): p. 2901–2913.

50. Giner-Lamia, J., L. López-Maury, and F.J. Florencio, CopM is a novel copper-binding protein involved in copper resistance in Synechocystis sp. PCC 6803. Microbiologyopen, 2015. 4(1): p. 167–85.

51. Imlay, J.A., The mismetallation of enzymes during oxidative stress. J Biol Chem, 2014. 289(41): p. 28121–8.

52. Ibrahim, I.M., S.J.L. Rowden, W.A. Cramer, C.J. Howe, and S. Puthiyaveetil, Thiol redox switches regulate the oligomeric state of cyanobacterial Rre1, RpaA and RpaB response regulators. FEBS Letters, 2022. 596(12): p. 1533–1543.

53. Kadowaki, T., Y. Nishiyama, T. Hisabori, and Y. Hihara, Identification of OmpR-family response regulators interacting with thioredoxin in the cyanobacterium Synechocystis sp. PCC 6803. PLoS One, 2015. 10(3): p. e0119107.

54. McKnight, B.M., S. Kang, T.H. Le, M. Fang, G. Carbonel, E. Rodriguez, S. Govindarajan, N. Albocher-Kedem, A.L. Tran, N.A.-O. Duncan, O. Amster-Choder, S.S. Golden, and S.A.-O. Cohen, Roles for the Synechococcus elongatus RNA-Binding Protein Rbp2 in Regulating the Circadian Clock. (1552-4531 (Electronic)).

55. Kobayashi, I., S. Watanabe, Y. Kanesaki, T. Shimada, H. Yoshikawa, and K. Tanaka, Conserved two-component Hik34-Rre1 module directly activates heat-stress inducible transcription of major chaperone and other genes in Synechococcus elongatus PCC 7942. Mol Microbiol, 2017. 104(2): p. 260–277.

56. Novichkov, P.S., A.E. Kazakov, D.A. Ravcheev, S.A. Leyn, G.Y. Kovaleva, R.A. Sutormin, M.D. Kazanov, W. Riehl, A.P. Arkin, I. Dubchak, and D.A. Rodionov, RegPrecise 3.0 – A resource for genome-scale exploration of transcriptional regulation in bacteria. BMC Genomics, 2013. 14(1): p. 745.

57. Giner-Lamia, J., L. López-Maury, J.C. Reyes, and F.J. Florencio, The CopRS Two-Component System Is Responsible for Resistance to Copper in the Cyanobacterium Synechocystis sp. PCC 6803. Plant Physiology, 2012. 159(4): p. 1806–1818.

58. Ostrowski, M., S. Mazard, S.G. Tetu, K. Phillippy, A. Johnson, B. Palenik, I.T. Paulsen, and D.J. Scanlan, PtrA is required for coordinate regulation of gene expression during phosphate stress in a marine Synechococcus. The ISME Journal, 2010. 4(7): p. 908–921.

59. Jiang, H.-B., W.-Y. Song, H.-M. Cheng, and B.-S. Qiu, The hypothetical protein Ycf46 is involved in regulation of CO2 utilization in the cyanobacterium Synechocystis sp. PCC 6803. Planta, 2015. 241(1): p. 145–155.

